# Distinct molecular mechanisms of stress habituation in the mouse hippocampus

**DOI:** 10.1101/2025.03.04.641433

**Authors:** Rebecca Waag, Lukas von Ziegler, Oliver Sturman, Emanuel Sonder, Justine Leonardi, Selina Frei, Katharina Gapp, Pierre-Luc Germain, Johannes Bohacek

**Affiliations:** Laboratory of Molecular and Behavioral Neuroscience, Institute for Neuroscience, Department of Health Sciences and Technology, ETH Zurich, Switzerland; Neuroscience Center Zurich, ETH Zurich and University of Zurich, Switzerland; ETH Zurich 3R Hub, ETH Zurich, Switzerland; Laboratory of Epigenetics and Neuroendocrinology, Institute for Neuroscience, Department of Health Sciences and Technology, ETH Zurich, Switzerland; Computational Neurogenomics, Institute for Neuroscience, Department of Health Sciences and Technology, ETH Zürich, Zurich, Switzerland; Laboratory of Statistical Bioinformatics, Department for Molecular Life Sciences, University of Zürich, Zurich, Switzerland

## Abstract

Chronic stress is a risk factor for neuropsychiatric disorders, making the ability to adapt to repeated stress a crucial determinant of mental health. On a molecular level, it remains unclear whether repeated exposure to stress is characterized by habituation - a decreased responsiveness to the same stimulus - or by the emergence of new, adaptive responses. Here, we explore how the tightly regulated molecular response triggered by acute restraint stress becomes altered after repeated restraint exposure. Transcriptomic sampling of the mouse hippocampus at multiple time points revealed that repeated stress leads to widespread habituation, damping stress-induced gene expression of all stress-responsive genes. However, we find no evidence for the emergence of new response profiles or alterations in baseline gene expression. Using single-cell multi-omics, we show that these findings hold true across cell types, and we reveal cell type specific patterns of habituation. Transcriptomic and chromatin accessibility profiles identify two distinct mechanisms that contribute to the observed habituation patterns: an early cAMP-associated mechanism that is related to blunted transcription after chronic stress, and a late corticosterone-dependent mechanism that is linked to a shortened transcriptional response. These extensive data are integrated, along with our previous work, into an <u>interactive app</u>, providing a uniquely detailed molecular resource that characterizes the acute stress response and the process of habituation across the genome.

## Introduction

In moments of acute stress, our bodies rapidly mobilize energy resources to ensure survival and initiate a complex molecular response to meet increased metabolic demands (Floriou-Servou et al., 2021). These widespread molecular changes resemble a symphony orchestrated by numerous neurotransmitters, peptides and hormones (Joëls and Baram, 2009; Floriou-Servou et al., 2021). Temporally, the response can be broadly divided into fast and slow components. The fast component, occurring within milliseconds, is mediated by the release of neurotransmitters and peptides like glutamate, GABA, corticotropin releasing hormone (CRH), noradrenaline or dopamine. The slow component is mediated primarily by the hypothalamus-pituitary-adrenal (HPA) axis, with corticosterone (CORT) reaching the brain only several minutes after stress initiation (Kitchener et al., 2004; Revest et al., 2005; Thoeringer et al., 2007; Droste et al., 2008; Qian et al., 2011). At the transcriptional level, where our molecular understanding of the acute stress response is most detailed, gene expression unfolds in waves and concludes within several hours after stress initiation (Reul, 2014; von Ziegler et al., 2022). By manipulating individual signaling pathways, studies begin to reveal which stress-mediators contribute to these stress-induced changes (Revest et al., 2005, 2014; Gerstner et al., 2022; Privitera et al., 2024).

To prevent wear-and-tear, exhaustion, and the onset of stress-related disorders, the stress response must adapt when stress occurs repeatedly and becomes chronic (McEwen and Akil, 2020; Nestler and Russo, 2024). However, adaptation to stress can come in many flavors. First, repeated stress exposure can lead to habituation, the reduced responsiveness to a repeated stimulus (Thompson and Spencer, 1966; Grissom and Bhatnagar, 2009). Habituation has been described, particularly for the HPA axis, across a variety of different stressors (Armario et al., 1984; Campmany et al., 1996; Bhatnagar et al., 2002; Babb et al., 2014). On a molecular level, stress-induced expression of immediate early genes (IEGs, like *Fos* or *Arc*) also typically habituates after chronic stress exposure (Melia et al., 1994; Chen and Herbert, 1995; Martinez et al., 1998; Stamp and Herbert, 1999; Girotti et al., 2006). Second, stress sensitization can occur when repeated presentation of the same stressor, or different stressors, triggers a more intense response than the first presentation. This process is often associated with heightened arousal in post-traumatic stress disorder, or with increased cortisol release from hypertrophied adrenal glands (Konarska et al., 1990; Ulrich-Lai et al., 2006; Anisman, 2011; Ursin, 2014; Belda et al., 2015). Third, chronic stress exposure can lead to the emergence of novel molecular and circuit adaptations that differ qualitatively from acute stress exposures, effects that have been studied particularly in the context of stress resilience (Nestler and Russo, 2024).

On a genome-wide level, it remains unknown how the acute stress response itself adapts after repeated stress exposure, and which mechanisms might be driving it. Most of the work on the transcriptomic consequences of chronic stress has focused on long-term alterations that manifest in the brain after extended periods of chronic stress (Peña et al., 2019). Large gene expression screens have revealed widespread changes in stress-sensitive regions like ventral hippocampus (vHC), prefrontal cortex, hypothalamus or nucleus accumbens, but tissue was collected long after the termination of the last stressor (Bagot et al., 2016, 2017; Nasca et al., 2017; Brivio et al., 2023). To understand how the transcriptomic response to the stressor itself adapts when the stressor is presented repeatedly, we chose a widely used restraint stress model and dynamically profiled multi-omic changes across multiple time points after the first and subsequent stress exposures. We describe, in both sexes, how the transcriptional response to acute restraint stress unfolds over time. We show profound transcriptional habituation and blunted chromatin remodeling across different cell-types as the stressor becomes chronic, and we identify mechanisms that drive different temporal components of stress habituation.

## Results

### Transcriptomic dynamics after acute restraint stress in the hippocampus

Acute restraint stress (ARS) is a commonly used stress model, which activates the hypothalamus-pituitary-adrenal (HPA) axis and leads to strong transcriptional responses in various brain regions (Cullinan et al., 1995; Girotti et al., 2006). Because the dynamic molecular changes triggered by ARS have never been characterized in detail, we exposed adult male and female mice to 1h30 of restraint stress, and collected the ventral hippocampus (vHC) for bulk RNA-sequencing at baseline, 45min, 1h30, 3h, 5h30 and 24h after stress initiation (**Figure 1A**). Consistent with previously described transcriptional responses to various stressors (Floriou-Servou et al., 2018), we detected strong gene expression changes 45min after stress initiation, changes that evolve in waves and resolve within several hours (**Figure 1B**). While some differentially expressed genes were still detected 5h30 after stress initiation, none were detected 24h afterwards (**Figure 1D-E**). These dynamics and the overall return to baseline is also visible in a multi-dimensional scaling representation of the gene expression response (**Figure 1C, left**). To check whether the effects at 5h30 represent lingering transcripts from earlier transcription, we looked at unspliced reads as a proxy for active transcription. We no longer observed significant group differences at 5h30, suggesting that transcription had indeed returned to baseline at this point (**Figure 1C, right**). This is in line with our previous work showing that 4hrs after the termination of swim stress, the transcriptional response had returned to baseline (note that 5h30 after ARS *initiation* corresponds to 4hr after ARS *termination*) (von Ziegler et al., 2022). Because swim and restraint stress are two very different acute stressors in terms of duration, modality and physical activity, we compared the two transcriptional responses directly (**Figure 1F**). Although log2 fold-changes (logFC) were generally stronger in response to swim stress, and some genes peaked at different time points (**Figure 1G**), in terms of gene activation and suppression patterns, the two responses were extremely similar (**Figure 1H**) and highly correlated (**Figure 1I**). This corroborates our previous observation that acutely stressful challenges trigger a highly conserved transcriptional response in both sexes (von Ziegler et al., 2022; Privitera et al., 2024), even when profiled dynamically across time.

**Figure 1.**
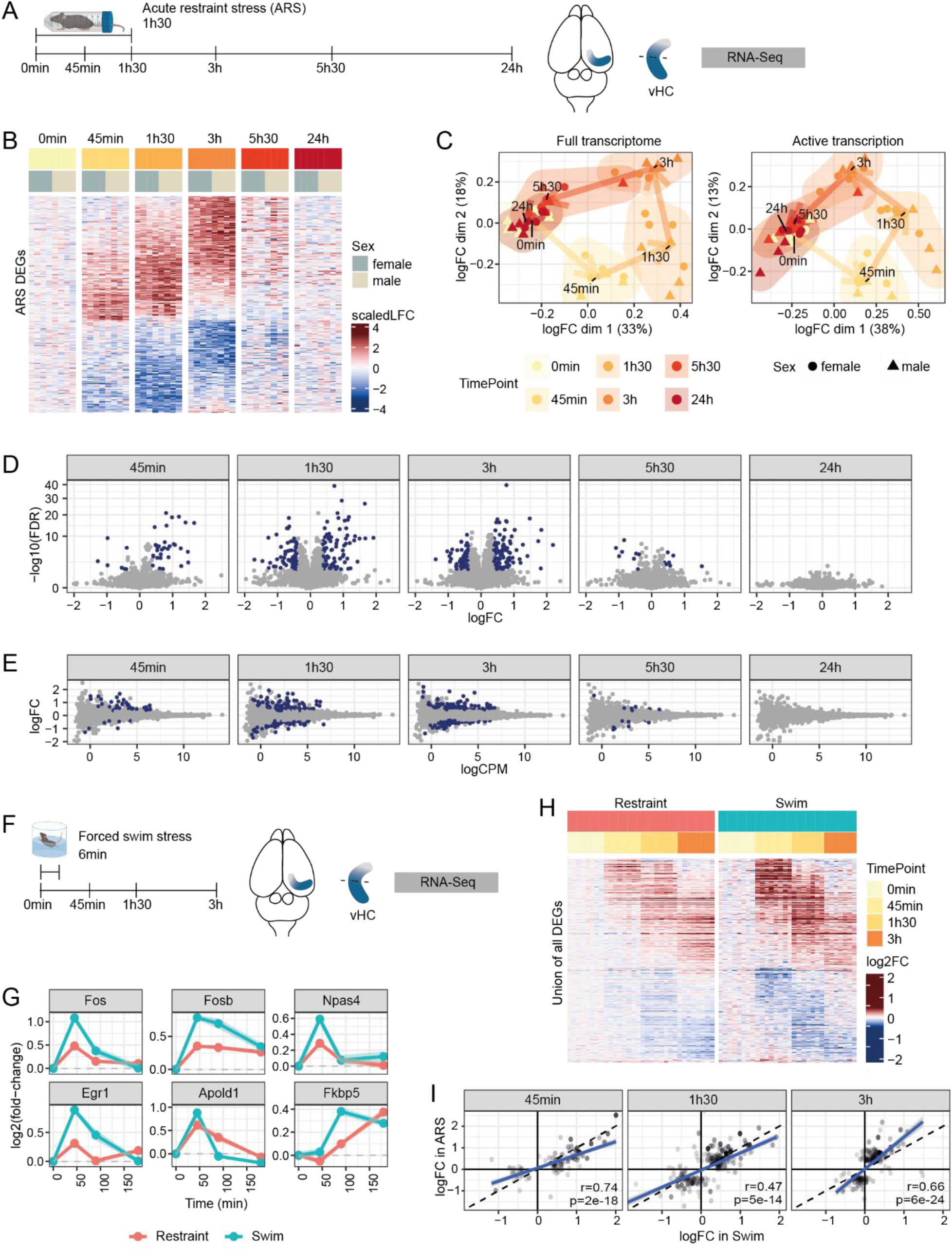
Dynamic transcriptional profiling of acute restraint stress. **A:** Design of the time-course profiling of the transcriptomic response to ARS. **B:** Relative gene expression of all genes affected by ARS. **C:** Leading logFC multi-dimensional scaling of the stress-responsive genes for the full transcriptome (left) and the active transcription (unspliced reads, right). **D:** Volcano plots showing significant (blue) differential expression at each time point following ARS. **E:** MA-plots showing significant (blue) differential expression at each time point following ARS. **F:** Design of the FSS experiments from von Ziegler et al. **G:** Expression, across the two datasets, of the union of genes significantly responding to either ARS or FSS (only common time points shown). **H:** Expression profile of important stress-responsive genes across both paradigms. The shaded area represents the standard deviation across animals. **I:** Comparison of the logFC at each common time-point of the union of genes significantly responding to either ARS or FSS. The dashed line is the diagonal, while the blue line represents a linear fit of the data.

### Global transcriptomic damping in response to repeated stress

To examine how the molecular response to ARS changes after repeated stress exposure, we subjected male and female mice to chronic restraint stress (CRS). CRS is one of the most widely used chronic stress models, it leads to habituation and results in long-term behavioral, molecular, and structural changes in the hippocampus and other brain regions (McEwen, 1999). As the transition from acute to chronic stress is hard to define, we exposed mice to 10 or 20 days of daily 1h30 restraint stress. The vHC was collected for bulk RNA-sequencing at baseline, 45min, 1h30, 3h, 5h30 and 24h after the last stress initiation (**Figure 2A**). Due to large cohorts, the experiments had to be conducted separately. Thus, additional handling control groups were added to both the 10d and 20d CRS experiments, for which tissue was collected at baseline, 45min and 24h after an ARS exposure. This allowed us to compare groups across cohorts, but it also enabled detailed within-cohort comparisons. Each group contained 8 mice, and each mouse served as a biological replicate (192 mice in total, a detailed schematic of the experimental design is provided in **Supplementary Figure 1**).

**Figure 2.**
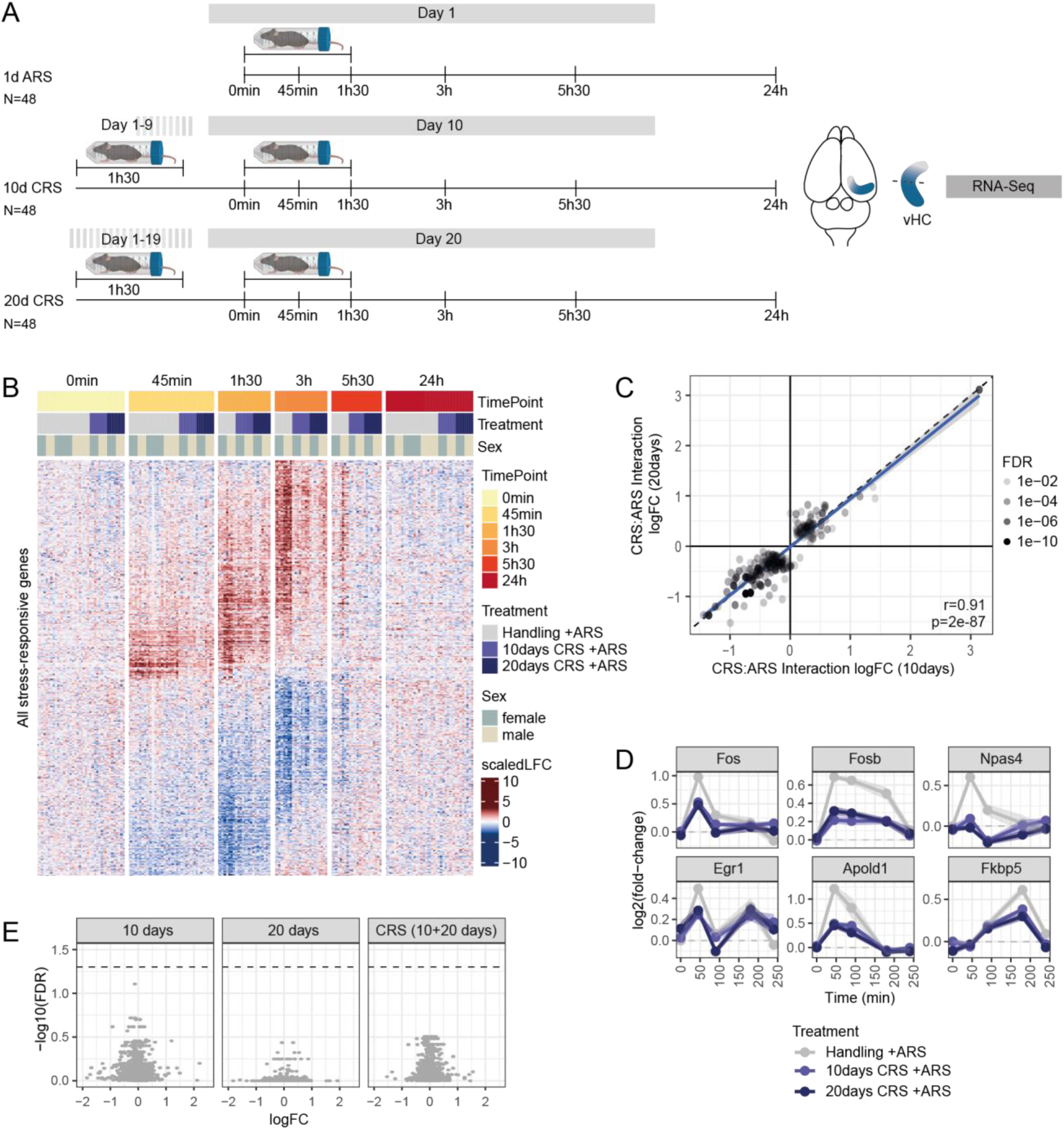
Global transcriptomic damping in response to repeated restraint stress. **A:** Design of the CRS experiment. **B:** Relative gene expression of all stress-responsive genes showing a general damping of the response. **C:** The lack of a global deviation in CRS:stress interaction coefficients across the two treatment durations indicates that overall, the two treatment durations elicit a similar damping. **D:** Key example genes showing similarly damped response across the two treatments. **E:** Volcano plot of the differential expression at baseline between CRS and handling samples, showing no significant changes. The dashed line indicates a 0.05 FDR threshold.

Across both sexes and treatment durations (10 and 20 days of CRS), we observed a profound damping of the ARS response in repeatedly stressed animals. Virtually all stress-responsive genes showed a reduced response in at least one time-point in CRS animals (**Figure 2B**). Surprisingly, we did not observe any overall difference in the magnitude of damping between 10 and 20 days of CRS, as indicated by a strong correlation between the CRS:ARS interaction fold-changes, which captures how each CRS treatment impacts the response to ARS (**Figure 2C** and **Supplementary Figure 2A**). This reveals that, at the level of gene expression, 10 days of CRS are sufficient to elicit strong transcriptional habituation of the acute stress response. **Figure 2D** shows the change in expression of prominent stress-responsive genes across the 3 treatment groups. Finally, we could not observe any significant expression differences between CRS and handling animals at baseline (**Figure 2E**). Although surprising given the extensive literature reporting long-term changes following chronic stress exposure (McEwen, 1999), this indicates that the habituation after CRS is not the result of a change in baseline gene expression. Thus, we next explored which mechanisms could account for the damping of stress-induced gene expression.

### Different mechanisms explain the temporal dynamics of habituation

The profiling of transcriptional changes over multiple time points allowed us to follow the temporal dynamics of stress habituation in unprecedented detail. To quantify these dynamics, we used the angle on the centered multidimensional scaling plot as a pseudo-temporal labeling of the samples across the global stress response (**Figure 3A**). This revealed that, in comparison to the first stress exposure, the global transcriptional response in CRS animals (10 or 20 days) was reduced at 45min.

**Figure 3.**
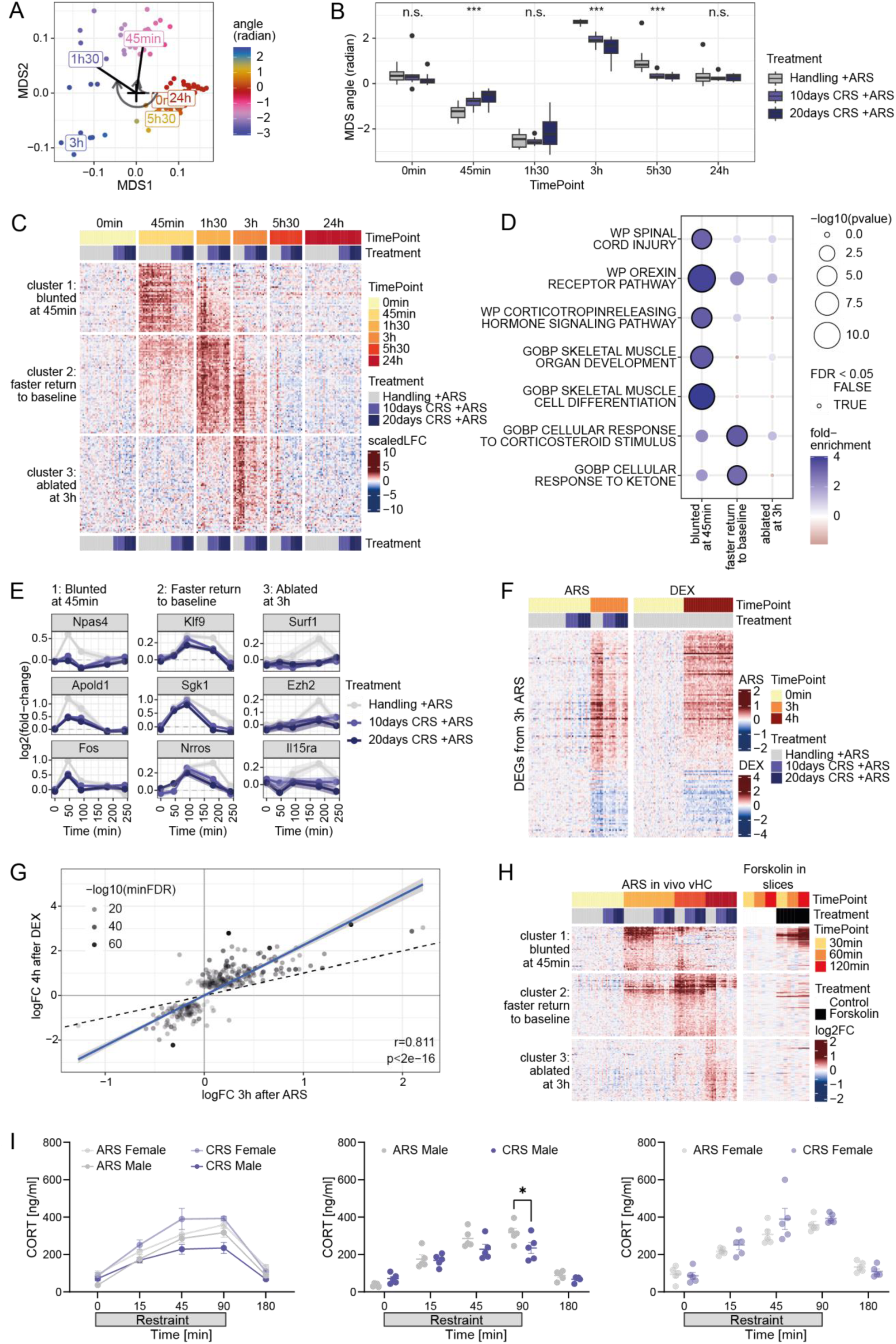
Temporal dynamics reveal different mechanisms of habituation. **A:** The angle on the multidimensional scaling (MDS) plot provides a pseudo-time labeling of the samples (shown are the animals without chronic stress). **B:** Using the MDS angle of each sample as pseudo-time, one can observe a significant damping of the early response in CRS animals, as well as a more rapid return to baseline. *** indicates adjusted p-value<0.001 (both treatments pooled for testing). **C:** Main patterns of damping across upregulated genes. **D:** Gene Ontology / Wikipathway over-representation analysis across the damping clusters. **E:** Example genes from each damping cluster. **F:** A comparison with DEX IP injection from Gerstner et al., 2022 shows that the changes observed at 3h ARS are recapitulated by DEX IP injection. **G:** Scatterplot of the logFC of the union of genes changed after 4h DEX IP or 3h ARS. **H:** Expression of the damping clusters in hippocampal slices upon forskolin-induced LTP (Chen et al., 2017) recapitulates the upregulated of the genes blunted at 45min, but not of the other clusters, as expected given the absence of GR signaling. **I:** CORT measures after ARS or 10d CRS in male and female mice. 3-way ANOVA (Treatment*TimePoint*Sex) reveals strong sex effect in CORT response (left, ****p<0.0001). Males and females are tested separately using a 2-way ANOVA (Treatment*TimePoint) with Sidak’s multiple comparison (middle, right). Data expressed as mean ± SEM. *adjusted p-value<0.05.

At 1h30 the molecular response peaked similarly in all groups, but returned to baseline much faster in CRS animals (**Figure 3B**). To further explore these different temporal dynamics, we used spectral clustering to capture groups of genes with similar transcriptional dynamics across time (**Supplementary Figure 2B**). Having previously shown that most transcriptional downregulation following acute stress is explained by RNA half-lives rather than direct transcriptional control (von Ziegler et al., 2022), we restricted the clustering to upregulated genes (see downregulated in **Supplementary Figure 2B** and **E**). We resolved three main damping patterns (**Figure 3C**): Cluster 1 captured transcripts that were blunted already at 45min. Cluster 2 captured transcripts that peaked normally in CRS animals, but returned to baseline faster. Cluster 3 captured transcripts that were normally upregulated at 3h but failed to do so in CRS animals. Enrichment analyses comparing the genes in each cluster to all expressed genes or to all stress responsive genes revealed converging results (**Figure 3D** and **Supplementary Fig 2D**): The genes blunted already at 45min (cluster 1) were especially enriched for CRH and Orexin receptor signaling pathways, as well as GO terms related to the vasculature (**Figure 3D**). Genes in this cluster particularly included neuronal IEGs, such as *Fos*, but also early response genes from the vasculature such as *Apold1* (**Figure 3E, left**). Instead, genes that peaked normally at 1h30 but returned to baseline faster in CRS animals (cluster 2) were enriched for response to abiotic stimulus and oxygen levels, as well as response to corticosteroids (**Figure 3D**). In line with this, one of the most specific glucocorticoid-responsive genes, *Fkbp5* (Zannas et al., 2016; Häusl et al., 2021), was normally upregulated at 1h30, but significantly damped at later time points (**Figure 2D**), and the glucocorticoid regulated kinase *Sgk1* showed a similar pattern (**Figure 3E, center**). The genes ablated at 3h (cluster 3) did not show enrichment that passed multiple testing correction, yet some exemplary genes are shown (**Figure 3E, right panel)**. Testing whether the three clusters showed enrichment for curated gene targets of transcription factors (based on the collecTRI regulons (Müller-Dott et al., 2023)) yielded corroborating results (**Supplementary Figure 2C**): Genes in cluster 1 were most prominently enriched for targets of the cAMP response element binding CREB1 (2.3-fold, FDR~5e-4), as well as other factors including the Serum Response Factor (SRF). In contrast, genes from cluster 2 were most strongly enriched for targets of the glucocorticoid receptor (GR) (3-fold, FDR~1e-7), with no target enrichment for cluster 3. To further confirm the involvement of GR signaling, we compared the response to ARS with the hippocampal gene expression response to an IP injection of the GR agonist dexamethasone (DEX) (Gerstner et al., 2022). The vast majority of genes upregulated 3h after ARS (which are all damped in CRS animals, whether in cluster 2 or 3) were also upregulated by DEX injection (**Figure 3F-G**). Conversely, we had previously shown that in vitro treatment of CA1/CA3 hippocampal slices with the cAMP activator forskolin (Chen et al., 2017) recapitulates, in the absence of corticosterone signaling, many of the early gene expression changes seen in response to acute stress exposure (Floriou-Servou et al., 2021). Indeed, we replicate this in the current dataset, where most of the genes blunted at 45min (cluster 1) were upregulated by forskolin in vitro (**Figure 3H**). In contrast, this was true only for a small minority of those genes that return faster to baseline (cluster 2), and only for 1 gene from those genes with ablated expression at 3h (cluster 3). In light of these observations, we suggest that there are two mechanisms, a rapid, GR-independent early damping that is related to cAMP signaling, and a slower, GR-dependent damping that emerges at later time points. Interestingly, this holds true for both 10 and 20 days of restraint stress.

### Behavioral and neuroendocrine outcomes of habituation

We were surprised that 10 days of restraint stress led to strong molecular habituation that was indistinguishable on the whole transcriptome level from 20 days of restraint. Thus, we asked whether behavioral changes, typically reported after 3 weeks of restraint stress exposure (Nasca et al., 2017), are also apparent after 10 days. Therefore, we compared the behavioral profiles of control, ARS and CRS male and female mice in the open-field test 45min after the end of their first ARS exposure, or after 10 or 20 days of CRS exposure (**Supplementary Fig. 3A+B**). In general, we observe an increase in distance moved and supported rearing upon both 10d and 20d CRS, indicating an increase in general activity compared to control animals (**Supplementary Fig. 3C+E**), with male and female animals generally responding similarly (**Supplementary Fig. 3D+F**). Additionally, we find a trend in CRS animals spending less time in the center of the arena. This suggests that behavioral changes emerge 10 days after stress exposure, which are similar to the changes observed after 20 days of stress exposure. Given the similar behavioral response, and to increase power, we combined the animals of both experiments. This global analysis shows strong differences between those animals being stressed for the first time and those being repeatedly stressed. After CRS, mice are hyperactive (more movement and more supported rearing) and spend less time in the center, suggesting increased anxiety (**Supplementary Fig. 3G**).

Given that habituation of the HPA axis has been well-documented, and because we see a clear involvement in GR in the damping of certain clusters of genes (**Figure 3C**), we measured corticosterone (CORT) levels in the blood in response to an ARS challenge presented either for the first time (ARS) or after 10 days of restraint stress (CRS-ARS). In line with previous work (Kitay, 1961), ARS triggered a stronger CORT increase in females than males (**Figure 3I**). CORT levels increased also in the CRS-ARS condition, with a modest blunting of the CORT response in males, but not in females. This is surprising, considering that the molecular habituation is very strong in males and females (**Figures 2+3**), and given that the behavioral changes are very similar in both sexes (**Supplementary Figure 3**). In males, CORT levels showed a similar rise, but a lower peak response at 1h30. Thus, changes in CORT release cannot explain the molecular habituation in females, and they likely cannot fully explain the strong molecular habituation observed in males either. Therefore, we next explored whether differences in chromatin accessibility can explain the blunted transcriptional response after CRS.

### Whole-tissue chromatin accessibility does not explain molecular habituation

To test whether changes in chromatin prevent stress-induced activation of gene expression after repeated stress exposure, we performed ATAC-seq of male and female mice that were either handled or stressed for 10 days both at baseline and 45min after ARS initiation (**Figure 4A**). Notably, analyses were performed in the same animals that were used for RNA sequencing, by using the contralateral hemisphere of the ventral hippocampus. We chose the 45min time point, because earlier work had shown that within an hour after stress exposure clear chromatin remodeling can be observed (Caradonna et al., 2022). Indeed, we observe differences (particularly increases) in peak accessibility, upon ARS (**Figure 4B-C**). A differential promoter accessibility analysis yielded several genes with increased accessibility upon ARS (**Figure 4D**), including known stress-responsive genes such as *Fkbp5* or *Nfkbia*, and novel genes like *Phyhd1* (**Figure 4D-E**). We correlated changes in promoter accessibility upon ARS with changes in RNA, using the union of genes that were significant in either ATAC-seq or RNA-seq. Since RNA changes persist much longer than changes in active transcription, and DNA accessibility relates to the latter, we focused on unspliced RNA at 45min, and observed a weak, but significant correlation (**Figure 4F, left**). Of note, some genes (e.g. *Hif3a* and *Fkbp5*) show an increase in promoter accessibility already at 45min, while the effect on gene expression can only be observed at later time points, indicating that chromatin accessibility foreshadows changes in transcription. We therefore performed the same comparison with the transcriptome at the next time point (1h30), and observed a slightly higher correlation (**Figure 4F, right**).

**Figure 4.**
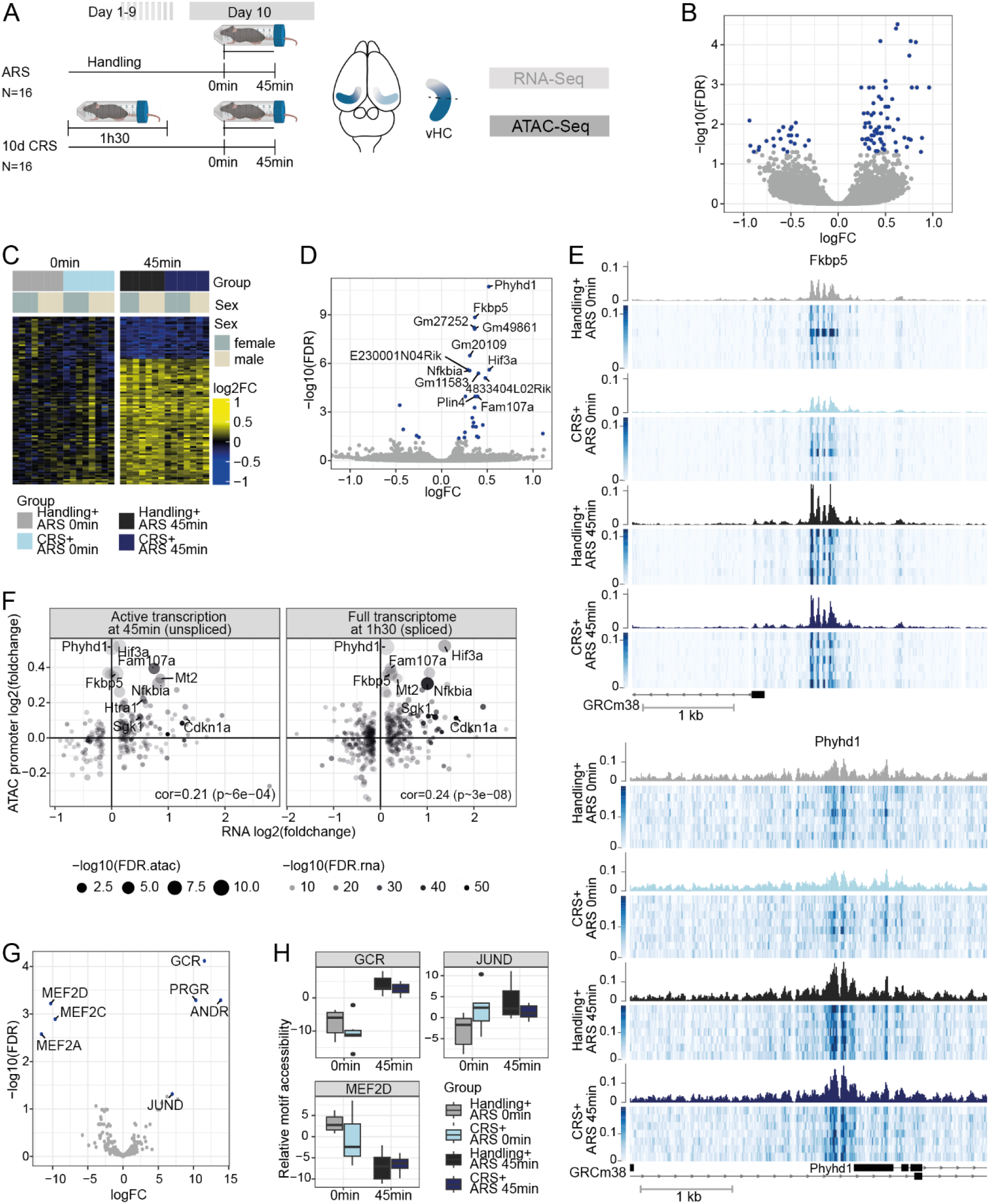
Chromatin accessibility in response to acute and repeated stress. **A:** Experimental design of the CRS experiment. **B:** Volcano plot of the differentially-accessible peaks upon 45min ARS (peaks with FDR<0.05 are in blue). **C:** Heatmap of the relative accessibility in statistically significant peaks. **D:** Volcano plot of the differential promoter (2kb upstream the TSS) accessibility 45min after ARS. Significant (FDR<0.05) genes are in blue, and the top ones are labeled. **E:** Tn5 insertion coverage across samples in the two promoters most significantly affected by ARS at 45min, *Fkbp5* (top) and *Phyhd1* (bottom). **F:** Volcano plot of the differential motif accessibility upon ARS (labeled are those with FDR<0.05). **G:** Relative motif accessibility of the most relevant representative of each of the three families of significant motifs. **H:** Correlation between changes in RNA and changes in the corresponding promoter accessibility upon stress. In the left panel, the change in promoter accessibility 45min after ARS is correlated to the change in active transcription (unspliced reads) at the same time point, while in the right panel it is correlated to changes in the full transcriptome at the next time point (1h30).

We next searched for transcription factors whose putative binding sites changed in accessibility (**Figure 4G-H**). The most significant motif was of the glucocorticoid receptor (as well as the very similar motifs of other hormone receptors), which showed an increase in accessibility, followed by MEF2-related motifs, showing decreased accessibility. Finally, we also observed a weaker, but significant increase in accessibility of the JUND motif (**Figure 4G-H**).

While we could identify major changes in accessibility upon ARS, we could not detect any change associated with CRS, either at baseline or in interaction with ARS. This means that 10 days of CRS do not alter ARS-induced chromatin remodeling. Even when focusing on the genes damped at 45min or 1h30, we could not observe any general pattern in the accessibility of their promoter in response to CRS. Thus, chromatin accessibility does not explain the damping in gene expression after repeated stress exposure. However, accessibility changes occur faster and are more transient than RNA changes, thus it is possible that we have missed CRS-associated changes in accessibility and TF activity that have resolved at the 45min time point. Additionally, it is possible that bulk ATAC-seq lacks the sensitivity to detect CRS-related changes, especially if they happen in a subset of cells. To address these two caveats, we turned to single-nuclei analysis of both transcriptome and chromatin accessibility across several time points.

### Transcriptomic single-cell dynamics of the acute stress response

It has recently been shown that stress-induced changes have cell-type specific effects, and that the analysis of bulk tissue can obscure biologically meaningful changes (Miranda et al., 2023; Waag and Bohacek, 2023). To test how habituation to the acute stress response evolves over time across cell types, we exposed female mice to ARS after 10 days of handling (handling-ARS), or after 10 days of chronic restraint stress (CRS-ARS), and collected the vHC at 15min, 45min and 3h after initiation of the last stress exposure (n=3 mice per group) (**Figure 5A**). We performed simultaneous single-nucleus RNA-seq and ATAC-seq on the same nuclei (10X Multiome). We sequenced a total of 215349 nuclei across 23 samples, and after stringent quality filtering (see methods) we retained ~160k cells with a median of 7343 UMIs per cell. Non-neuronal cells included oligodendrocytes and oligodendrocyte progenitor cells (OPC), with a fraction of committed oligodendrocyte progenitors in-between (**Figure 5B**). Microglia and astrocytes had prominent clusters, and we could also identify perivascular macrophages (PVM). Although lower in number, we could observe clear populations of pericytes, endothelial cells, smooth muscle cells, and some smaller subpopulations of vascular cells with unclear signatures, and we merged them as ‘vascular’ cells for downstream analysis.

**Figure 5:**
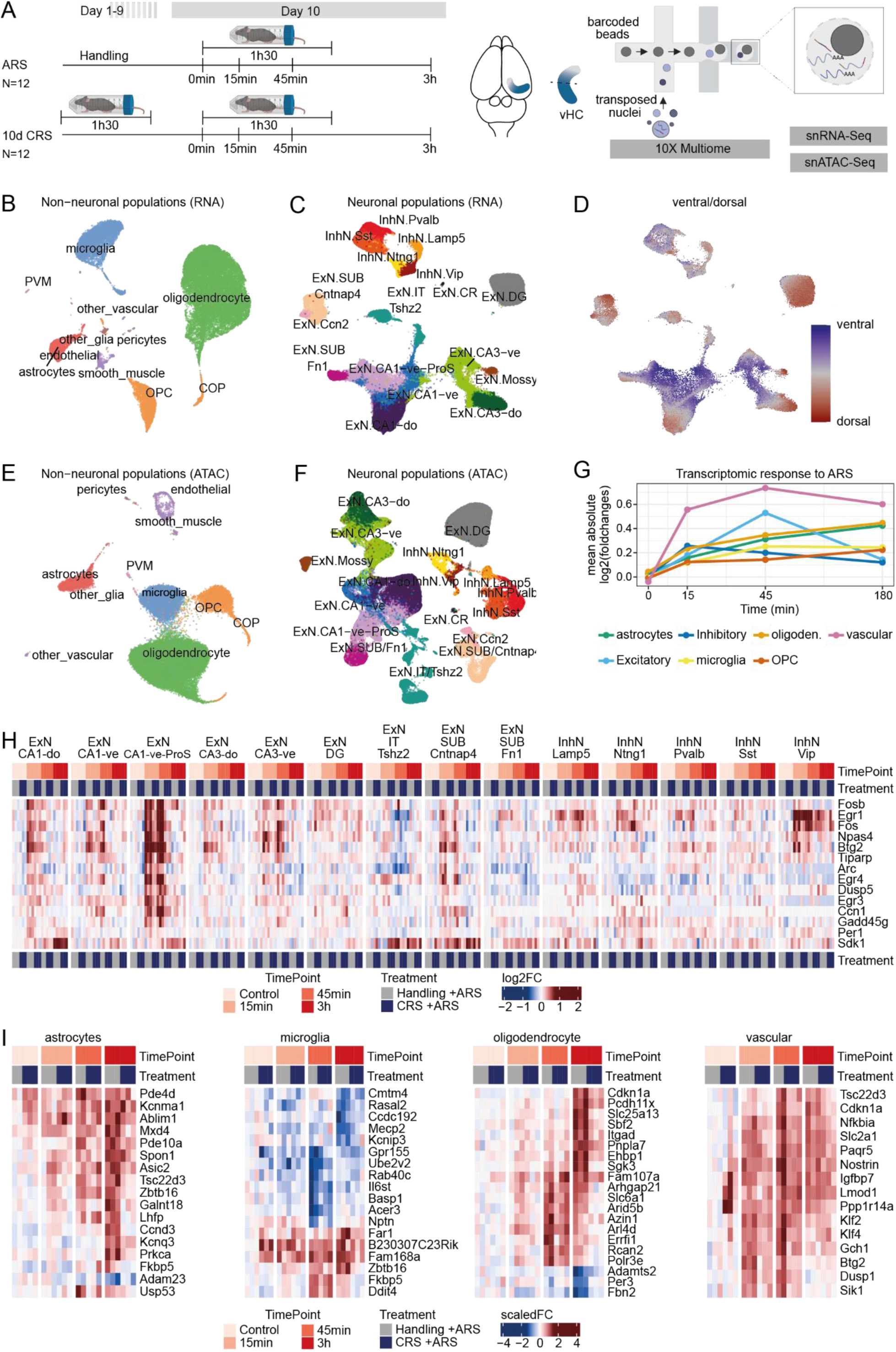
The transcriptomic response to acute stress and habituation across cell types in the ventral hippocampus. **A:** Experimental design of the CRS experiment and the multiomic pipeline. **B:** UMAP of the neuronal cell populations in the transcriptomic space, colored and labeled by cluster. **C:** UMAP of the neuronal populations in the ATAC space, showing a good segregation of the cell clusters identified from the RNA modality. **D:** UMAP of the neuronal populations in the RNA space, in which the cells are colored by their Ventral-dorsal score based on the CA1/3 signature from Cembrowski et al. (elife 2018). Note that the signature is based on cells from the pyramidal layer, and might not be meaningful for other cellular populations (e.g. inhibitory neurons). **E:** UMAP of the non-neuronal populations in the transcriptomic space. **F:** UMAP of the non-neuronal populations in the ATAC space, coloring cells by their cluster from the RNA modality. **G:** Overview of the global, dynamic response to ARS across broad cell classes. **H:** Pseudo-bulk expression of genes found significantly changing upon ARS in more than one neuronal cell type. **I:** Pseudo-bulk expression of the genes significantly changing upon ARS in each broad class of non-neuronal cells (max 20 genes shown per cell type).

The different neuronal subpopulations are shown in **Figure 5C** (marker expression is shown in **Supplementary Figure 4A**). Inhibitory neurons clustered by known type (Sst, Pvalb, Vip, Ntng1 interneurons, and Lamp5 neurogliaform interneurons), while excitatory neurons clustered by region, first with a clear distinction between granule cells from the dentate gyrus, mossy cells from the hilus, and CA1 and CA3 excitatory neurons. Within the pyramidal layer (i.e. CA1-3), cells then clustered by relatively more dorsal vs ventral regions (**Figure 5C-D**) based on signatures from Cembrowski et al. (Cembrowski et al., 2018). Dorsal scores were generally lower, and fewer cells had a high dorsal score, in line with the fact that only ventral hippocampi were analyzed (**Supplementary Figure 4B**). This shows that the ventral-dorsal gene expression gradient is preserved – and can be resolved – even within the ventral portion. In addition, we identified a gradient of cells spanning from the CA1 to the pro-subiculum and the distal subiculum (Fn1+), as well as a separate population most likely from the posterior subiculum (with *Ly6g6e* and *Cntnap4* expression). Another gradient especially characterized by high *Camk2d* expression extended from the ventral CA1 population. Based on correlations with the Allen brain atlas, they appear to be neurons from the hippocampus-tegmental amygdala (HATA) and/or hippocampus-inferior temporal cortex interfaces. These populations were the only ones to (mildly) vary in abundance across samples, consistent with their location on the border of the dissected tissue. Of note, the transcriptome-based cell type labels were also clearly segregated in the ATAC-seq embedding, both for non-neuronal (**Figure 5E**) and neuronal (**Figure 5F**) populations, corroborating the robustness of the cell type assignment.

To date, no dynamic profiling of the acute stress response on the single-cell level exists in the hippocampus. Therefore, we first examined the transcriptional changes to ARS on a pseudo-bulk level (**Supplementary Figure 5**). Globally, the different broad cell types all responded to ARS early on (**Figure 5G**). In neuronal cell types, the response peaked at 15min (inhibitory neurons) or 45min (excitatory neurons), before strongly decreasing at 3h. In contrast, the response of all non-neuronal cell types was sustained at 3h. In neurons, a particularly strong response was observed in CA1 pro-subiculum and VIP interneurons, which upregulated a number of major IEGs already 15min after stress initiation, such as *Fos*/*Fosb*, *Npas4*, and *Btg2* (**Figure 5H**). With the exception of vascular cell types, which also responded very quickly, other non-neuronal populations were slower to respond (**Figure 5I)**. The significant genes in non-neuronal populations included a number of prominent stress-induced genes, such as *Apold1* in vascular cells (Fan et al., 2023), the cell cycle regulator *Cdnkn1a*, or the GR target *Fkbp5* across multiple cell types. Of note, some changes, such as the strong upregulation of the cAMP/cGMP-degrading enzyme *Pde10a* specifically in astrocytes upon stress (**Figure 5I** and **Supplementary Fig 5**), had not been detected in the bulk transcriptome, presumably because the gene is also highly expressed in other glial populations at baseline, underlining the value of single-cell profiling.

### Stress habituation occurs across cell-types

Visual inspection of the heatmaps in **Figure 5H-I** suggests that across cell types, the response to ARS was generally damped after CRS, with different temporal patterns emerging for different cell types. To investigate this systematically, we tested, within each cell type, which stress-induced gene expression profiles were altered by CRS at any time point. Several genes were significantly affected by CRS, with a clear pattern of ARS-responsive genes habituating in CRS animals (**Figure 6A**). This was also the case for several prominent stress-responsive genes (**Figure 6B**). Notably, while *Npas4* upregulation appeared to be completely ablated in CRS animals at 45min based on the bulk transcriptomic data, the single-cell data revealed an upregulation at 15min also in CRS animals across multiple excitatory neuron subtypes (**Supplementary Figure 6B**). However, in CRS animals this upregulation was weaker, and had already returned to baseline at 45min across cell types, while it remained upregulated in animals who experienced the stressor for the first time (**Figure 6B** and **Supplementary Figure 6B**), consistent with the bulk data. In contrast to our bulk sequencing data, a number of genes appeared differentially expressed at baseline in CRS animals (**Figure 6C**). While some of these candidates showed a variable expression pattern across ARS time-points (e.g. *Cacna1e* and *Vwf*), some looked more coherent, such as *Pde1a* in VIP interneurons, a gene that is involved in cGMP/cAMP homeostasis (Delhaye and Bardoni, 2021). To ascertain these findings, we also specifically tested for differences at baseline, which revealed some statistically significant candidates (**Supplementary Figure 6A**). While the identification of rare, cell-type specific baseline gene expression changes reveals the power of single-cell sequencing, these limited changes cannot explain the widespread damping of stress-induced activity across cell types. Next, we leveraged the single-cell resolution to explore stress-activated cells in more depth.

**Figure 6:**
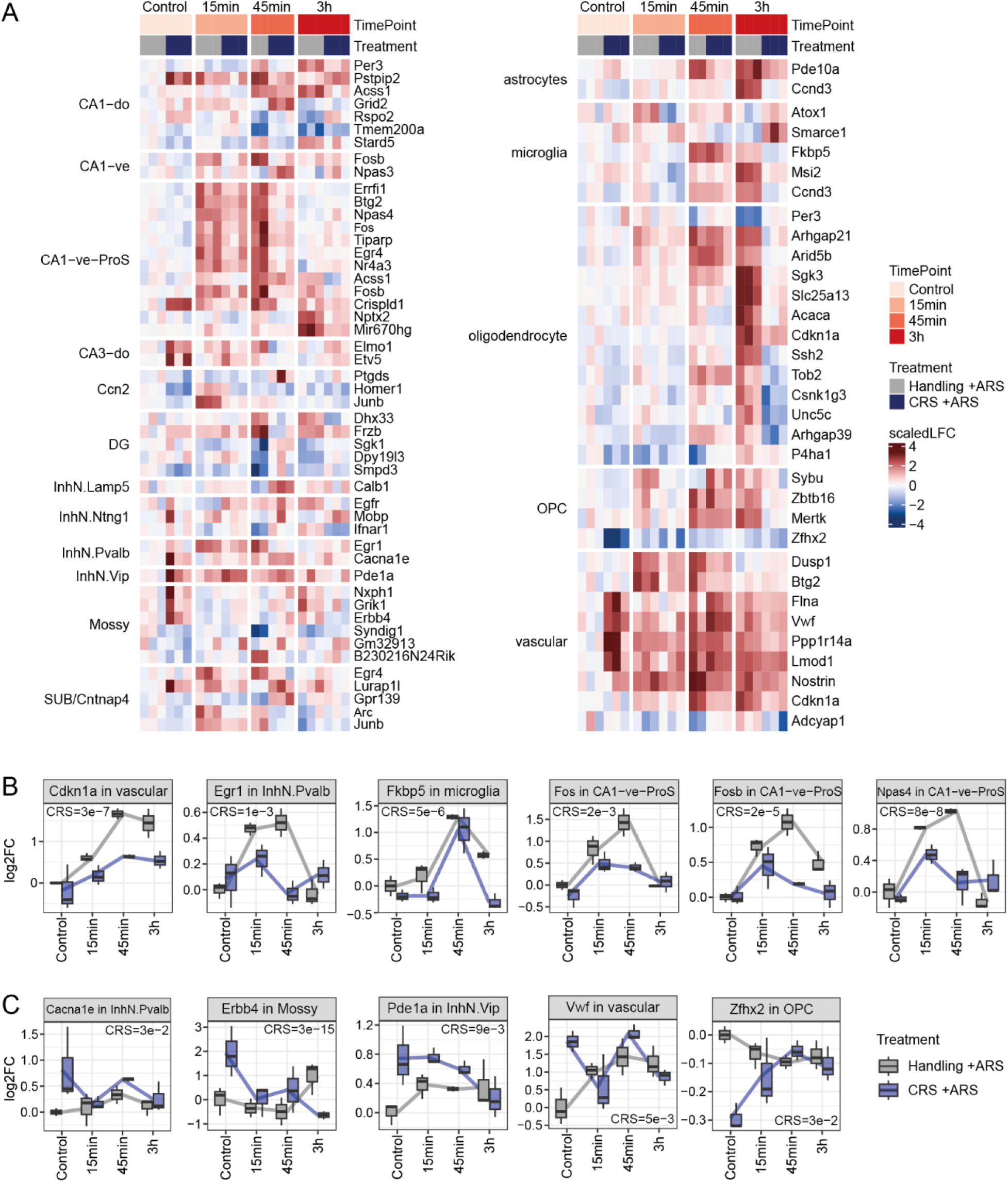
Transcriptomic stress habituation occurs across cell types. **A:** Relative expression of genes showing a significant impact of CRS on the stress response across cell types. **B:** Key IEGs showing a significant habituation in CRS; shown are the cell types in which they are the most significant. **C:** Candidate genes with a significant effect of CRS at baseline. Reported significance values are from a likelihood ratio test comparing the full model (ARS time points and CRS covariates, i.e. ~ARS*CRS) to a null model without CRS covariates (CRS p-value), and adjusting for multiple testing using Benjamini & Hochberg’s method. A significant CRS q-value means that CRS affects either the baseline gene expression or that the gene responds differently to ARS after CRS.

### After habituation, fewer cells respond to stress

A dampened transcriptomic response to stress could be due to fewer cells responding, or to the same proportion of cells having a weaker response. cFOS or ARC stainings for instance indicate that only a fraction of hippocampal neurons respond to salient, stressful events (Cullinan et al., 1995). Leveraging the power of single-cell transcriptomics, we estimated the recent activity of single neurons using an activity-dependent transcription (ADT) score based on genes that are differentially expressed upon ARS at the pseudo-bulk level (see methods). The ADT score indicates, for each cell, whether it looks more like a cell that does not respond to ARS (score of 0) or like the most stress-responsive cells (score of 1). As expected, while only a minority of the cells showed ADT, these were unequally spread across subpopulations (**Figure 7A**), with the highest activity in subiculum, pro-subiculum and CA1 neurons. Using an ADT score above 0.5 as a threshold for an activated neuron, we observed that the proportion of activated neurons doubled upon the first ARS exposure, a response that was blunted after CRS (**Figure 7B**). In contrast, while the median ADT score of activated neurons did show a significant increase upon ARS, the difference was very small in magnitude (roughly 5%, **Figure 7C**). Together, this indicates that most of the damping is due to fewer neurons being activated, rather than the same number of neurons being activated less strongly.

**Figure 7:**
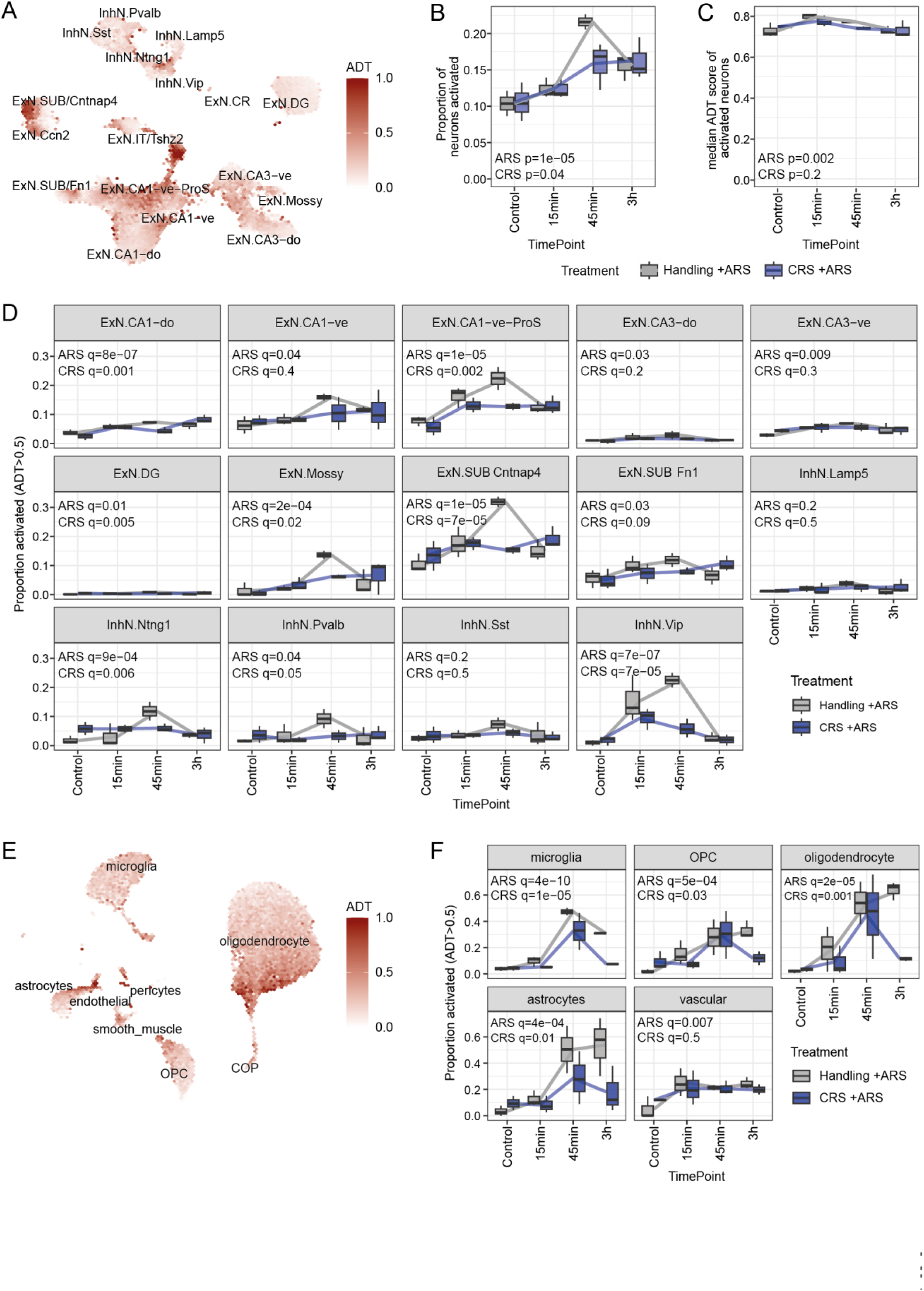
Activity-dependent transcription across populations. **A:** UMAP of neuronal populations, colored by their activity-dependent transcription (ADT) score. **B-C:** Proportion of neurons activated (**B**) and median ADT score or activated neurons (**C**) across samples, indicating a decreased number of recruited neurons upon stress in CRS animals. **D:** Proportion of activated neurons per pseudo-bulk sample across neuronal subpopulations, using the mean of the ADT and neuroestimator scores. **E:** UMAP of non-neuronal populations, colored by their ADT score. **F:** Proportion of activated cells across broad non-neuronal cell types. Reported p-values are from a likelihood ratio test, comparing the full model (ARS time points and CRS covariates, i.e. ~ARS*CRS) to a null model without the time points (ARS p-value) or without the CRS covariates (CRS p-value). A significant CRS q-value means that CRS affects either the baseline activation or that activation in response to ARS differs after CRS.

We next repeated this analysis at the level of individual cell types, which showed that the habituation after CRS is cell-type specific, with the strongest blunting of ADT observed in VIP interneurons and in several excitatory cell populations (**Supplementary Figure 7**). To corroborate these results, we also used an independent method to estimate neuronal activation from single-cell data, neuroestimator (Bahl et al., 2024). Although the estimated baseline activation levels across subpopulations varied depending on the methods, the main patterns upon ARS and CRS were largely reproduced for inhibitory cell types, with less congruence in excitatory populations (**Supplementary Figure 7**). **Figure 7D** shows the proportions of each cell type activated based on the average of the single-cell scores from the two methods. Consistently, VIP interneurons showed the highest change in stress-induced activity. Strong responses in ADT were also observed in CA1 and subiculum populations, in mossy cells, as well as in other inhibitory neuron subtypes (Pvalb and Ntng1). Notably, the temporal dynamics varied across cell types, e.g. VIP interneurons were strongly activated already at 15min, while other inhibitory neurons showed a more delayed activation. In all these populations, the strong ARS-induced change was blunted or completely absent after CRS, especially at 45min (**Figure 7D**).

Contrary to the neuroestimator method, our ADT approach is applicable to any cell type, and we thus used it to estimate the proportion of cells recruited across non-neuronal populations (**Figure 7E**). Only a minority of vascular cells reacted to ARS, whereas the majority of astrocytes and oligodendrocytes strongly reacted upon first ARS exposure, with microglia and OPC showing intermediate activation. In all glial cell types, but not in vascular cells, the ARS-induced activation was blunted in CRS animals, especially at 3h. This reduction was particularly strong for astrocytes, dropping roughly by half already from 45min on. In contrast, oligodendrocytes seem to be recruited similarly in response to stress, but while their activity is sustained across 3 hours in handling animals, activity drops sharply at this time point in CRS mice.

### Stress-associated changes in chromatin accessibility across cell types

To investigate the mechanisms underlying the observed transcriptional changes following acute stress, and the blunting after chronic stress exposure, we next turned to the changes in chromatin accessibility across cell types. In line with the bulk ATAC-seq data, we identified several transcription factors (TFs) whose motifs were differentially accessible upon ARS in many cell types (**Figure 8A** and **Supplementary Figure 8**). Across stress-responsive neuronal cell types (i.e. especially CA1, pro-subiculum and VIP neurons), we could observe a strong increase in the accessibility of Fos/Jund motifs at all time points following ARS (**Figure 8A-B**). However, this was not well coupled to changes in the accessibility of cAMP-response elements (CRE): while the Creb1 motif appeared to increase in accessibility at 15min in Ntng1 and VIP interneurons, this was not the case in in most excitatory neurons, where it instead showed a trend in the other direction (**Figure 8A-B**). This was especially surprising given the clear recruitment of these neurons and the known rapid CREB-dependent transcription upon depolarization (Sheng et al., 1991; West et al., 2002). To confirm this finding, we inferred CREB activity with an independent method (fastMLM, see methods) based on experimental binding sites in neurons (Loupe et al., 2024). This corroborated the previous patterns of increased CREB activity in VIP neurons 15min after ARS, and a general decrease (especially at 45min) in stress-responsive excitatory neurons (**Supplementary Figure 9A**). To confirm the expected involvement of CREB in this paradigm, we tested motif enrichment within regions accessible in neurons in proximity to genes that respond to stress rapidly (i.e. genes that are significantly increased by stress at 15min in single-nuclei data or at 45min in the spliced bulk transcriptome). We identified several over-represented motifs, with CREB1 showing the strongest enrichment (**Supplementary Figure 9B**; motifs from the KLF family were also highly enriched). A similar analysis using experimental binding data highlighted a number of factors, including EGR1 and CREB1 (**Supplementary Figure 9C**), and CREB1 was the only factor significant in both analyses (**Supplementary Figure 9D**). One possible explanation for these results is that the expected cAMP chromatin response following depolarization is already over at 15min, and the observed reduction in CRE accessibility represents the shutdown of this response. If this were the case, then the increased CRE accessibility visible e.g. in VIP neurons at 15min could instead reflect a later (or sustained longer) activation. In line with this, we also observed a strong increase in EGR1 motif accessibility in VIP neurons at later time points (**Figure 8B**), and a strong and prolonged increase in *Egr1* expression in this cell type after stress exposure (**Figure 5H**). While *Egr1* is highly expressed across most excitatory neuronal populations, and in most cases also upregulated upon stress, the fold-changes are relatively modest in comparison to its 21-fold upregulation in VIP interneurons (**Supplementary Figure 9E**). Interestingly, EGR1 motif accessibility in VIP neurons is blunted in VIP neurons after CRS (**Figure 8B**), and stress-induced *Egr1* expression returns to baseline much faster after CRS exposure (**Supplementary Figure 9E**). Although the role of *Egr1* in stress habituation and synaptic plasticity has been well-described (Duclot and Kabbaj, 2017), the strong involvement of VIP neurons has thus far not been reported.

**Figure 8.**
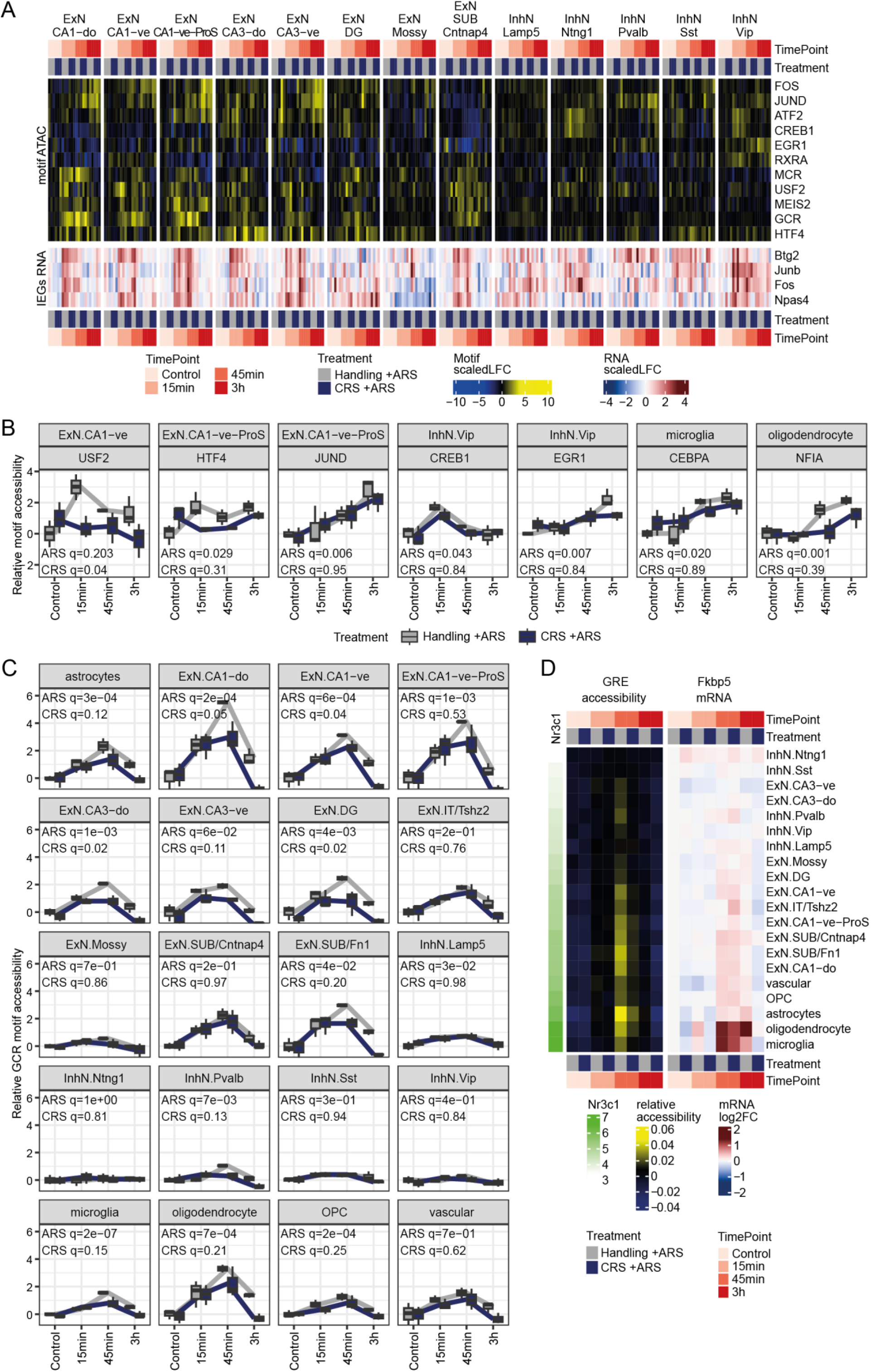
Stress-induced chromatin remodeling across time and cell types. **A:** A selection of top motifs responding in accessibility to ARS across neuronal cell types. To relate these to RNA, the expression of the top 4 IEGs is shown below in the corresponding samples. **B:** Relative motif accessibility of selected motifs. **C:** GR motif accessibility across all cell types. Reported significance values are from moderated F-statistics, comparing the full model (ARS time points and CRS covariates, i.e. ~ARS*CRS) to a null model without the time points (ARS p-value) or without the CRS covariates (CRS p-value). Adjustment for multiple testing using Benjamini & Hochberg’s method. A significant CRS p-value means that CRS affects either the baseline motif accessibility or that the motif accessibility responds differently to ARS. **D:** Overview of the relationship between GR (Nr3c1) expression (left in green), relative changes in accessibility of Glucocorticoid Response Elements (GRE, center), and Fkbp5 mRNA expression (right) across cell types and treatment.

Across all cell types, the strongest and most significant ARS-induced changes in motif accessibility consistently came from the glucocorticoid receptor and highly similar steroid hormone receptor motifs (**Figure 8A,C** and **Supplementary Figure 8**), with increased accessibility already at 15min and extending to later time points (**Figure 8C**). Notably, the increase in GR motif accessibility upon ARS was much weaker in interneurons, which also did not upregulate its target *Fkbp5*. We hypothesized that this could be due to differences in expression of the GR gene (*Nr3c1*), and indeed observed that, across cell types, the magnitude of the increase in GR motif accessibility in response to ARS was proportional to the expression of *Nr3c1*r (**Supplementary Figure 9G**). Although *Nr3c1* has a lower expression in inhibitory neurons, it is nevertheless relatively high, suggesting their lower glucocorticoid sensitivity involves other mechanisms. In addition, we identified several other TF motifs that strongly respond to ARS in non-neuronal cell types (**Supplementary Figure 8**), in particular CEBPA in microglia and NFIA in oligodendrocytes (**Figure 8B**).

### Cell-type specific chromatin accessibility yields mechanisms for habituation

Finally, we analyzed how ARS-induced DNA accessibility changed after a history of CRS. We found little evidence of habituation in motif activity among neuronal populations. Notable exceptions are the glucocorticoid receptor, the MAPK-dependent and Fos-interacting gene USF2, as well as HTF4/HEB, encoded by the *Tcf12* gene and reported to interact with the AP-1 complex at cAMP response elements (Di Rocco et al., 1997; Wang et al., 2022) (**Figure 8A-B**). As none of these genes are differentially expressed in CRS animals, their effect would have to be either through interactors or post-transcriptional regulation. While the ARS-induced change in GR motif accessibility was similar between handling-ARS and CRS-ARS animals at 15min across all cell types, we observed clear damping in the CRS-ARS group at 45min in many cell types **(Figure 8C)**. After 3h, GR motif accessibility completely returned to baseline in the CRS-ARS group, but not in handling-ARS animals (**Figure 8C**). Notably, the increased GR accessibility already 15min after stress exposure is in contrast with the transcriptional impact of GR signaling, as glucocorticoid-responsive genes showed transcriptional changes only at later time points. We illustrate this with *Fkbp5*, one of the most specific GR target genes; *Fkbp5* expression was only upregulated at the RNA level at 45min (across several cell types), and expression was significantly damped in many cell types 3hrs after stress initiation (**Figure 8D**, **Supplementary Figure 9F**). Altogether, these results suggest that glucocorticoids reach the brain and trigger GR translocation to the nucleus within 15min after initiation of stress, foreshadowing GR-dependent transcription at later time points. Again, this supports the hypothesis that the transcriptional damping already observed at 15min is GR-independent, and that GR signaling is likely responsible for much of the damped response at later time points.

## Discussion

Here, we explored the adaptation of the acute stress response on the level of gene expression and chromatin accessibility in the mouse hippocampus. While previous work had extensively profiled both the molecular changes induced by acute stress (Floriou-Servou et al., 2021; von Ziegler et al., 2022), and the long-lasting changes after chronic stress (McEwen, 2000, 2017; Bagot et al., 2016; Nasca et al., 2017; Peña et al., 2019), it was unknown how the acute stress response adapts on a molecular level as a stressor becomes chronic. By profiling the acute stress response at multiple time points and after different durations of CRS, we show strong habituation of the stress response across the vast majority of stress-responsive genes, albeit with different temporal dynamics. Based on these dynamics, we can dissociate a blunting of immediate early genes from a damped and shortened response of glucocorticoid-sensitive genes.

### Early GR-independent transcriptional damping

Within minutes of an acute stress exposure, the transcriptional machinery becomes activated and gene expression changes can be observed (Revest et al., 2005; Roszkowski et al., 2016). These changes are pioneered by well-known IEGs, such as *Fos*, *Npas4* and *Apold1,* which group together in cluster 1 (**Figure 3C**). *Fos* is known to strongly habituate to CRS on the transcript and protein level in various brain regions (Melia et al., 1994; Chen and Herbert, 1995; Martinez et al., 1998; Stamp and Herbert, 1999). Both our bulk and snRNA-Seq results replicate *Fos* habituation at 45min across cell types. Similarly, we find that virtually all stress-responsive IEGs show habituation at least at one time point. We observe that the members of cluster 1 are strongly enriched for CREB1 and SRF transcription factor activity, both crucial for IEG induction (Bahrami and Drabløs, 2016). Indeed, members of cluster 1 are upregulated following forskolin treatment of hippocampal slices (Chen et al., 2017) (**Figure 3H**), but not in response to Dex, thus indicating that cAMP signaling is at least partially involved. Importantly, this correlation is not found for clusters 2 and 3, suggesting that upregulation of the targets in these clusters is largely cAMP-independent. Our single-nucleus ATAC-seq analysis confirms the involvement of CREB-dependent signaling, showing that stress-induced IEGs are enriched for the CREB1 motif in close proximity peaks that show increased accessibility.

What remains unclear, however, is whether after habituation some IEGs are blunted from the start, or whether their initial response is preserved but transcription shuts down faster. *Npas4,* for example, shows complete habituation 45min after repeated stress in our bulk RNA-Seq, but we still find a significant – albeit weak – increase in *Npas4* transcripts at 15min in several cell types in our single-nucleus data. In a few cell types, however, *Npas4* seems to be completely silenced across time points (**Supplementary Figure 6B**). Due to the lack of a finer temporal resolution within the first 15min of stress initiation, we cannot fully resolve the question whether IEG transcription is similarly initiated between ARS and CRS. We can, however, resolve one critical question based on our new method for quantifying activity dependent transcription (ADT): We find that, across cell types, the blunted response to ARS after a history of CRS is due to fewer cells being activated, rather than the same proportion of cells showing a weaker activation.

### GR-dependent damping

Acute restraint stress strongly induces HPA axis activation. While CORT levels in the blood rise within a few minutes, CORT is detected in the brain after 10 min (Thoeringer et al., 2007; Droste et al., 2008). On the bulk and single-cell level, our transcriptomic analyses reveal genes that peak similarly at 1h30 after stress onset, but shut down faster and return to baseline levels at the 3h time point in animals with a history of chronic stress, while still being strongly upregulated after the first ARS exposure. Converging evidence shows that these transcripts are enriched for known GR target genes, such as *Fkbp5* (Zannas et al., 2016; Häusl et al., 2021), and they overlap with DEX-responsive genes from an independent analysis. In both our bulk and single-cell ATAC sequencing, the strongest ARS-induced changes were associated with an increase of GR motif accessibility, consistent with previous research (Caradonna et al., 2022). After CRS exposure, this ARS-induced increase is largely damped at 45min and 3h across cell types. However, we do observe a strikingly similar increase in GR accessibility at 15min (**Figure 8C**), thus indicating that indeed GR activation is initiated similarly to ARS, but shortened. CORT levels measured in the blood support the notion that the initial rise is similar in animals with or without a history of prior stress. However, at later time points, CORT dynamics paint a more complex picture, suggesting that CORT levels habituate slightly in males, but not in females. Notably, several reports find that the CORT levels habituate strongly in response to repeated restraint stress (Campmany et al., 1996; Bhatnagar et al., 2002; Babb et al., 2014), yet not all studies observe this habituation (Makino et al., 1995; Umemoto et al., 1997) and we suspect that varying the time of day (unpredictable restraint stress) might have prevented HPA-axis habituation in our setup. While our results support previous reports of sex differences in HPA-axis regulation and a more pronounced CORT response in females (Kitay, 1961; Galea et al., 1997; Bohacek et al., 2015), they contrast with the fact that we observe no corresponding sex differences in overall stress habituation patterns. These data suggest that in males, a blunting of the HPA-axis response could account – at least in part – for the damping of GR-sensitive genes after repeated stress exposure, whereas in females processes downstream of CORT release seem to be at play. This could be related to known sex differences in glucocorticoid binding globulin (Galea et al., 1997). More work is needed to reveal the mechanisms behind these unexpected effects.

### Is habituation an adaptive or maladaptive process?

It remains unclear whether the profound transcriptomic stress habituation we observe is beneficial to the organism, or whether it represents a maladaptive change. Habituation is often conceived as a strategy to decrease the impact of the repeated activation of the metabolically demanding stress response (McEwen, 1998). Additionally, the failure to habituate has been associated with the development of cognitive and stress-related disorders, such as PTSD (Orr et al., 1995; de Tommaso et al., 2014; McDiarmid et al., 2017; Cavanagh et al., 2018). However, our behavioral data suggest that the molecular habituation, which we observe on the level of every single animal after 10 and 20 days of CRS, is accompanied by a consistent decrease in time spent in the center of the OFT arena, a behavior considered to indicate increased anxiety (Prut and Belzung, 2003). Other findings using different behavior tests after CRS point in the same direction, e.g. decreased time spent in the open arm of the EPM (Kim and Han, 2006; Chiba et al., 2012), a decrease in sucrose preference (Chiba et al., 2012; Mao et al., 2022) as well as increased time floating in the forced swim test (Kim and Han, 2006; Chiba et al., 2012; Nasca et al., 2017). These phenotypes are associated with increased anxiety-like behavior, anhedonia, and behavioral despair (Gencturk and Unal, 2024), suggesting that the reduced or more efficient response to repeated stress comes at the cost of negative emotional changes. Indeed, recent work has shown that blunted corticosterone responses cause behavioral traits akin to PTSD (Seah et al., 2022; Monari et al., 2024). Notably, the molecular habituation cannot be explained by a baseline shift, as we detect – across large numbers of samples – no differences in baseline gene expression levels between handled and chronically stressed animals. This underscores that the molecular habituation can only be revealed when an acute stress challenge is imposed, and when the stress-response is profiled dynamically across time. As we lack a brain-wide understanding of transcriptional habituation to repeated stress, it remains unclear how brain regions other than the hippocampus contribute to the emerging behavioral phenotypes.

### Limitations

As the acute stress response represents a dynamic system, and because the border between acute to chronic stress is blurry, we faced two challenges: 1) To capture temporal dynamics we had to collect samples at many time points across several hours; 2) We had to capture the gradual onset of adaptive changes over days to weeks. Regarding the former, our data suggests that habituation of GR-dependent transcription is due to a more transient transcriptional activation, rather than a blunted increase in the first place. For GR-independent stress habituation that occurs much earlier after stress exposure, our single-nucleus data suggest that in some cell types the activation of IEGs might be completely abolished. However, we would need more time points during the first 15min after stress initiation to firmly corroborate this claim. Regarding the gradual emergence of habituation over time we find that both 10 and 20 days of CRS result in a very consistent behavioral phenotype as measured in the open field. Additionally, we do not find a difference in the transcriptional stress response between animals being exposed to 10 or 20 days of CRS. This indicates that the actual habituation process already happens earlier in time. Indeed, on the physiological level, a clear reduction of the stress-induced CORT response can already be observed upon the second or third stress exposure in some stress models (De Boer et al., 1990; Babb et al., 2014), but whether this holds true for transcriptional blunting remains to be tested. Importantly, our study focuses on group effects and does not account for individual variance in the ability to habituate. Although the transcriptional and behavioral habituation itself was highly consistent across animals, the extent of habituation likely varies among individuals, highlighting a potential area for future investigation. Finally, due to the requirement of a chromatin accessibility readout, we profiled single nuclei instead of single cells, which restricts the RNA readout to the nucleus and excludes transcripts located in the cytoplasm and neuropil. While this approach prevents dissociation-induced transcriptional artifacts (Waag and Bohacek, 2023), the resulting data is dominated by nascent transcripts (Bakken et al., 2018). This offers a clear snapshot of ongoing transcription, but lacks the cumulative representation of lingering transcripts seen in our bulk RNA data.

## Conclusion

Using multi-omic bulk and single-cell sequencing, we identify distinct mechanisms that contribute to the emergence of genome-wide molecular habituation to a recurrent stressor. We integrate these new data with our previously published sequencing analysis of the acute stress response (von Ziegler et al., 2022), thus creating a searchable, open-source database. These data offer a detailed characterization of the “healthy” stress response and lay the foundation to explore how this response might vary between individuals and how it may become altered in stress-related diseases.

## Material & Methods

### Animals

C57BL/6J (C57BL/6JRj) mice were obtained from Janvier (France). Mice were maintained in a temperature- and humidity-controlled facility on a 12 h reversed light–dark cycle (lights off at 09:30 a.m.) with food and water ad libitum. Male and female mice were housed in groups of 2-5 animals per cage and used for experiments when 2–3 months old. All tests were conducted during the animals’ active (dark) phase. All procedures were carried out in accordance with Swiss cantonal regulations for animal experimentation and were approved under licenses ZH106/2020, ZH067/2022 and ZH103/2023.

### Stress paradigm

Mice were exposed to restraint stress for 1h30 in a 50 ml Falcon tube with a large air hole. All stress exposures were carried out between 10:00 a.m. and 5:00 p.m. Animals were single-housed 24 hours prior to the final stress exposure to reduce confounding effects induced by cage mates on the day of the experiment (Bohacek et al., 2015). For acute stress experiments, animals were exposed to one restraint session (1h30). The restraint tube was placed in the homecage for the duration of stress. For chronic stress experiments, animals were subjected to either 10 or 20 consecutive days of daily restraint stress at varying times of the day. Control animals and animals exposed to acute stress were gently handled daily for an equal number of days. Gentle handling involved lifting the animals by the tail, simulating the handling method used before restraint stress.

### CORT measurement

To assess CORT levels, blood samples were collected via tail prick at 0min, 15min, 45min, 90min and 180min after the onset of restraint. Blood samples were stored at 4°C overnight to allow for clotting and centrifuged at 2500 g for 10 min at 4°C. The serum supernatant was collected and stored at −80 °C until further analysis.

Corticosterone levels in the serum were measured using an ELISA kit (Invitrogen, EIACORT) according to the manufacturer’s instructions. A 1.5 μl serum sample was diluted 1:100 for the assay. Absorbance was measured at 450 nm using a plate reader, and final concentrations were determined based on the provided standards.

### Tissue collection

At the indicated time point after initiation of stress, mice were euthanized by cervical dislocation and decapitation. When tissue was collected at 15min, 45min or 1h30 after stress initiation, mice were removed from the restraint tube at the indicated time point and immediately euthanized for brain collection. The brain was quickly dissected on a cool surface and isolated ventral hippocampi were snap-frozen in liquid nitrogen and stored at −80°C until further processing.

### Whole tissue RNA extraction

One snap-frozen ventral hippocampus per animal was used to obtain transcriptional profiles. Samples were processed in a multiple block design with every block containing one replicate of each condition and sex. Processing within blocks was randomized. Samples were homogenized in 500 µl Trizol (Invitrogen 15596018) using metal beads (Qiagen 69989) in a tissue lyser bead mill (Qiagen, Germany) at 4°C for 2 min at 20 Hz. RNA was extracted according to the manufacturer’s recommendations. RNA purity and quantity were determined with a UV/V spectrophotometer. RNA integrity was assessed with high sensitivity RNA screen tape on an Agilent Tapestation/Bioanalyzer, according to the manufacturer’s protocol.

### Whole-tissue library preparation and sequencing

For experiment 1 (see **Supplementary Figure 1**), library preparation and sequencing was performed at the Functional Genomics Center Zurich (FGCZ) of University of Zurich and ETH Zurich. The TruSeq stranded RNA kit (Illumina Inc.) was used according to the manufacturer’s protocol. For bulk sequencing library preparation, the TruSeq stranded RNA kit (Illumina Inc.) was used according to the manufacturer’s protocol. The mRNA was purified by polyA selection, chemically fragmented and transcribed into cDNA before adapter ligation. Single-end (100nt) sequencing was performed with HiSeq 4000. Samples within experiments were each run on one or multiple lanes and demultiplexed. A sequencing depth of ~20M reads per sample was obtained.

For experiments 2+3 (see **Supplementary Figure 1**), library preparation and sequencing was performed at Novogene, UK. Briefly, RNA samples were used for library preparation using NEBNext® Ultra RNA Library Prep Kit for Illumina®. Indices were included to multiplex multiple samples. Briefly, mRNA was purified from total RNA using poly-T oligo-attached magnetic beads. After fragmentation, the first strand cDNA was synthesized using random hexamer primers followed by the second strand cDNA synthesis. The library was ready after end repair, A-tailing, adapter ligation, and size selection. After amplification and purification, insert size of the library was validated on an Agilent 2100 and quantified using quantitative PCR (Q-PCR). Libraries were then sequenced on Illumina NovaSeq 6000 S4 flowcell with PE150 according to results from library quality control and expected data volume. Samples within experiments were each run on one or multiple lanes and demultiplexed. A sequencing depth of ~20M paired reads per sample was obtained.

### Bulk transcriptome analyses

Bulk RNA-seq was analyzed as described in von Ziegler et al., 2022. Wherever surrogate variable (SV) analysis (using sva 3.50.0, (Leek et al., 2012) identified more than 5 SVs, we restricted it to 5. To identify damping patterns, sva-corrected, unit-variance-scaled logFCs of differentially-expressed genes (DEGs) across samples from time points 45min to 3h were first extracted, and values ceiled between −5 to 5 to restrict outlier influences. Genes were then clustered using the Spectrum 1.1 package with k=10. The MDS was computed through the plotMDS function of the limma package version 3.58.1 using the sva-corrected expression of the acute stress DEGs across Handling samples, and the CRS samples were then projected onto this space. The MDS was recentered so that all time point medians were equidistant to the origin before calculating angles. Cluster enrichment analysis was done using Fisher’s exact test (followed by FDR correction) on the KEGG, WIKIPATHWAYS and GO genesets of effective sizes ranging from 5 to 1000 from the MSigDB (Castanza et al., 2023), accessed from the msigdbr R package version 7.5.1.

### Nuclei isolation for ATAC-Seq

Nuclei isolation was performed as previously described (Grandi et al., 2022). All steps were performed on ice. Briefly, one vHC was homogenized in lysis buffer (26 mM Sucrose, 30 mM KCl, 10 mM MgCl2, 20 mM Tricine-KOH (pH 7.8), 1 mM DTT, 0.5 mM Spermidine, 0.15 mM Spermine, 0.3% NP-40, Complete protease inhibitor in nuclease-free water) by 10 strokes with pestle A and 20 strokes with pestle B. The sample was filtered through a 70 μm Flowmi strainer and centrifuged for 5 min at 400 g at 4°C. Supernatant was removed and additional lysis buffer was added to resuspend the pellet. The sample was then mixed with 50% Iodixanol solution (30 mM KCl, 26 mM Sucrose, 10 mM MgCl2, 20mM Tricine-KOH (pH 7.8), 50% Iodixanol solution in nuclease-free water) and underlaid by 30% Iodixanol solution (50% Iodixanol solution in lysis buffer) and 40% Iodixanol (50% Iodixanol solution in lysis buffer) layers. The sample was centrifuged for 20 min at 3000 g at 4°C with brakes off. The nuclei band was collected and mixed with ATAC buffer (10 mM NaCl, 10 mMTris-HCl (pH7.5), 3 mM MgCl2, 0.1% Tween20 in nuclease-free water). The nuclei count was assessed and 50’000 nuclei were transferred to a new tube with ATAC buffer. The samples was centrifuged for 10 min at 500g at 4°C and supernatant was removed.

### Preparation of ATAC libraries

Nuclei were resuspended in 50 μl of transposition mix (25 μl 2× TD buffer, 2.5 μl transposase (Diagenode, C01070012), 16.5 μl PBS, 0.5 μl 1% digitonin, 0.5 μl 10% Tween-20, and 5 μl nuclease-free water). Transposition reactions were incubated at 37 °C for 30 min while shaking at 1,000 r.p.m.. Tagmented DNA was purified using the MinElute Reaction Cleanup Kit (Qiagen, 28204). Purified DNA was amplified using the NEBNext high-Fidelity 2X master mix (New England Biolabs, Ipswich, Massachusetts, United States, M0541) and Diagenode primers (Diagenode, Marlborough, Massachusetts, United States, C01011036) at 72 °C for 5 min, 98 °C for 30 s and 5 cycles of 98 °C for 10 s, 63 °C for 30 s, and 72 °C for 1 min. A fraction of the amplification product was used to determine the number of additional cycles using qPCR, resulting in a total of 8-9 amplification cycles for all samples. A double-sided size selection (0.5X and 1.3X) was performed to remove large and very small library fragments. Insert size of the library was validated and quantified on an Agilent 2100 Bioanalyzer. Libraries were then sequenced on Illumina NovaSeq 6000 S4 flowcell with PE150 according to results from library quality control and expected data volume aiming for 50 Million read pairs per sample. Sequencing was performed at Novogene, UK.

### Analysis of bulk ATAC sequencing

Reads were trimmed with trimmomatic and aligned to the GRCm38 genome using bowtie2 with --dovetail --no-mixed --no-discordant -I 15 -X 2000. Duplicates were marked using Picard. Peaks were called using Genrich 0.6.1 (https://github.com/jsh58/Genrich) with ATAC mode with a FDR threshold of 0.05, discarding PCR duplicates and excluding the mitochondria and ENCODE blacklisted regions. Quantification was performed by counting the number of insertions events (i.e. shifted fragment starts/ends) within peaks or promoter regions. Differential analysis was performed with edgeR glmFit using a model including Sex, Treatment, ARS time point, and a Treatment:TimePoint interaction. For promoter-based analysis, an additional surrogate variable based on the SVA package was included. Insertion coverage tracks were generated (and plotted) with the epiwraps package v0.99.102, using binWidth=10L, shift=c(4L,-5L), type=“ends”, extend=3L, and includeDuplicates=FALSE.

Motif accessibility analysis was performed using chromVAR. For this purpose, peaks were resized to 300bp and insertions were counted as described above. 2000 iterations were used, and the z-scores were quantile-normalized as recommended in (Gerbaldo et al., 2024). Differential analysis was performed with limma using 1 surrogate variable.

### Nuclei isolation for single-cell experiments

Nuclei isolation was performed similarly as for ATAC-Seq, with the addition of RNase inhibitors as previously described (Grandi et al., 2022). All steps were performed on ice. Briefly, one vHC was homogenized in lysis buffer (26 mM Sucrose, 30 mM KCl, 10 mM MgCl2, 20 mM Tricine-KOH (pH 7.8), 1 mM DTT, 0.5 mM Spermidine, 0.15 mM Spermine, 0.3% NP-40, Complete protease inhibitor, 0.2 U/μl RNAse inhibitor in nuclease-free water) by 10 strokes with pestle A and 20 strokes with pestle B. The sample was filtered through a 70 μm Flowmi strainer and centrifuged for 5 min at 400 g at 4°C. Supernatant was removed and additional lysis buffer was added to resuspend the pellet. The sample was then mixed with 50% Iodixanol solution (30 mM KCl, 26 mM Sucrose, 10 mM MgCl2, 20mM Tricine-KOH (pH 7.8), 50% Iodixanol solution in nuclease-free water) and underlaid by 30% Iodixanol solution (50% Iodixanol solution in lysis buffer) and 40% Iodixanol (50% Iodixanol solution in lysis buffer) layers. The sample was centrifuged for 20 min at 3000 g at 4°C with brakes off. The nuclei band was collected and mixed with ATAC buffer (10 mM NaCl, 10 mMTris-HCl (pH7.5), 3 mM MgCl2, 0.1% Tween20, 0.2 U/μl RNase inhibitor in nuclease-free water) before the nuclei count was assessed. The sample was centrifuged for 5 min at 500g at 4°C and supernatant was removed. Diluted nuclei buffer (0.1 M DTT, 1 U/μl RNase inhibitor, 1X Nuclei buffer (10X) in nuclease-free water) was added to achieve the desired nuclei concentration.

### Preparation of single-nucleus Multiome libraries

10X Multiome libraries were prepared following the Chromium Next GEM Single Cell Multiome ATAC + Gene Expression Reagent Kits User Guide (CG000338, Rev. F, 10X Genomics) with a targeted nuclei recovery of 10’000 nuclei (1 sample failed due to clogging). 7 cycles were performed for sample index PCR during ATAC library preparation. 6 cycles were used for cDNA amplification and a total of 13 cycles was performed for sample index PCR for gene expression library preparation. RNA integrity was assessed on an Agilent Tapestation, according to the manufacturer’s protocol. Samples of each library type were pooled. Both libraries (gene expressions and ATAC) were then sequenced on one Illumina X Plus 25B flow cell on individual lanes with an adjusted read configuration (150-10-24-150 bp). A sequencing depth of ~50’000 paired reads per nucleus was targeted. Library preparation and sequencing were performed at the Functional Genomics Center Zurich (FGCZ) of University of Zurich and ETH Zurich.

### Single-cell analysis

#### Pre-processing and filtering

Pre-processing was performed using cellranger-arc 2.0.2 using the optimized mm10 transcriptome as described previously (Pool et al., 2023). A RNA quantification using alevin-fry (He et al., 2022) was also performed to estimate splicing proportions. Decontamination was performed using cellbender 0.3.0 (Fleming et al., 2023) with custom initialization, and doublet identification with scDblFinder 1.19.0 (Germain et al., 2022). For the RNA modality, scDblFinder was run using standard parameters twice (i.e. with two seeds) and the scores averaged, the AMULET method was used on the ATAC modality. Cells were excluded as doublets if they were called as such using both scDblFinder runs, or if their mean doublet score was above 0.2 and their amulet q-value below 0.01. Only cells with at least 500 fragments (after decontamination), less than 60% contamination, less than 1% mitochondrial reads, more than 20% unspliced fragments (before decontamination) and less than 50% of the fragments assigned to the top 10% features were used for further analysis.

#### Single-nucleus RNA-seq analysis

Cell annotation was performed based on the RNA modality. A first rough Seurat 5.0.1 clustering (Hao et al., 2024) was performed to split neuronal and non-neuronal cells, which were then processed separately using Seurat.

For neurons, 3500 variable features were selected using the disp method. Prominent activity-dependent genes were removed to limit the influence of the activation status on cell annotation, and known markers were added (see repository for the exact list). Batch correction was performed using harmony 1.2.1 (Korsunsky et al., 2019). The first 30 dimensions of the harmony embedding were then used to identify the 50 shared nearest neighbors used for clustering at resolution 1.3 and UMAP projection. Cell clusters with a median proportion of unspliced reads below 50%, a median doublet score above 0.3, a median proportion of mitochondrial reads above 0.02, a median proportion of contamination above 0.3, or half of their markers being more highly expressed in empty droplets than in cells were removed as low-quality clusters. Clustering was then performed again on the cleaned object. The clusters were then compared to the Allen brain atlas annotation using SingleR (Aran et al., 2019) for the purpose of annotation, in combination with known markers. In addition, we used the CA1/3 marker genes from Cembrowski et al (Cembrowski et al., 2016) to distinguish ventral and more dorsal populations.

For non-neuronal cells, we first used geo-sketching (Hie et al., 2019) to 6000 cells as implemented in sketchR 1.1.2 to increase sensitivity to rare cell types. We took the union of the top 2000 variable features identified using the disp and vst methods, as well as the top 1000 most expressed features in the sketched data and known markers. We then performed PCA on the entire dataset based on these features followed by harmony. The first 20 dimensions of the harmony embedding were then used for clustering and UMAP projection. Cell clusters with a median proportion of unspliced reads below 35% or a median doublet score above 0.2 were removed, as well as clusters of cycling cells based on the expression of Top2a, Ccnb2, Cdk1 and Ccnf. The remaining clusters were annotated manually based on known markers.

For ADT-based analysis (below) and visualization, only cells with at least 1000 fragments were used. Pseudo-bulk differential analysis was performed using edgeR with a ~TimePoint*Treatment model. If the median abundance of the cluster across samples was higher than 50 cells, one surrogate variable estimated using the sva package was further included in the model. For each cell type, genes were filtered using filterByExpr with min.count=20, and only genes with an average log(CPM) above 1.5 were considered. We then harnessed our vast bulk RNA-seq dataset to improve power by estimated global FDR in the pseudobulk data with the IHW package v1.30.0 (10.1038/nmeth.3885), using the FDR from the bulk data as covariate. Specifically, we used the smallest FDR between the spliced and unspliced bulk data from dropping all experimental covariates (for baseline differences between CRS and Handling, the FDR from the baseline bulk comparison was instead used). Genes which were below filtering thresholds at the bulk level, and which consequently had no attached bulk FDR, were assigned a value of 2 to have their weights estimated separately. IHW was then applied independently for each contrast, across cell types of the same class (i.e. neuronal/non-neuronal). Finally, a minimum absolute logFC of 0.5 was used in addition to FDR to identify differentially-expressed genes.

#### ADT scoring

The Activity-Dependent Transcription (ADT) score was computed separately for broad cell classes (with neurons grouped as either excitatory or inhibitory). For each class, we first defined a set of stress-induced genes with global (i.e. across time points) FDR below 0.15 (0.05 for non-neurons), an absolute logFC above 0.2 and a log(CPM) above 1. Since the latest time point is chiefly due to GR, we additionally required the gene to have a FDR<0.15 at either 15min or 45min. For each gene, we then computed the weighted average of its pseudobulk logFC at 15min and 45min, weighted by 1-sqrt(FDR) in the respective time point. These logFC estimates then served as the expected activity logFC, on which each cell was regressed. Specifically, we used Gamma-Poisson regression (using the glmGamPoi package 1.17.3 with global overdispersion and no normalization (Ahlmann-Eltze and Huber, 2021) of the cell’s counts for those genes on two covariates: the scaled sum expression of the genes across all cells of that type, and the expected logFC. The coefficient assigned to the second covariate then represents a raw activity score, which is however strongly influenced by the proportion of cells of that type that are in an activated state. To address this issue, we then scale the score for each cell, setting 0 to the median score in baseline animals, and 1 to the 98th percentile of the score. An ADT score of 0.5 then represents the point where cells are more similar to the most strongly activated cells than to the median baseline cell. For neurons, an alternative activity scoring was performed using the neuroestimator package version 0.1 (Bahl et al., 2024), and to circumvent the biases of both methods, the two scores were averaged for each cell.

#### Single-nucleus ATAC-seq analysis

Using the RNA-based clusters, fragments were merged for neuronal and non-neuronal cell types and peaks were called separately using MACS2 version 2.2.9.1 (Zhang et al., 2008) via the CallPeaks function of Signac 1.13.0 version (Stuart et al., 2021). Downstream analysis was performed on pseudobulks of the RNA-based clusters obtained using muscat version 1.16.0 (Crowell et al., 2020). For motif accessibility analysis, peaks were resized to 500bp before running chromVAR on each cell type separately and performing differential analysis as discussed for bulk ATAC-seq. To avoid reporting the several sets of highly similar motifs showing a common pattern, we first applied hierarchical clustering of motifs using the Yule similarity of their matches across peaks, and cutting the dendrogram at a height of 0.5. For each cell type, we took the significant motifs from the same cluster and excluded the ones for which the corresponding gene had an expression below 2 logCPM in that cell type. We then reported the remaining ones in decreasing order of expression. To validate motif-based patterns using ChIP-seq binding data, we first used liftover to translate the human brainTF peaks to mouse coordinates, and then overlapped the lifted neuronal peaks on the single-cell ATAC peaks. As chromVAR is not applicable to such data, we then used the next best method, fastMLM (after smooth quantile normalization in GC bins), from Gerbaldo, Sonder et al. (Gerbaldo et al., 2024).

### Open field test

Open-field testing took place inside sound insulated, ventilated multi-conditioning chambers (TSE Systems Ltd, Germany). The open field arena (45 cm x 45 cm x 40 cm [L x W x H]) consisted of four transparent plexiglas walls and a light gray PVC floor. Animals were tested for 10 min under dim lighting (4 lux). Video recordings were collected and distance, time in center, supported rears and unsupported rears were analyzed.

### Pose estimation using DLC and behavior analyses

DeepLabCut 2.0.7 (DLC) (Nath et al., 2019; Lauer et al., 2022) was used to track 13 body points of each animal. Tracked points included nose, head center, neck, right ear, left ear, body center, body center left, body center right, left hip, right hip, tailbase, tail center and tail tip. The four corners of the OFT were tracked additionally to automatically detect the arena boundaries in each recording. X and Y coordinates of DLC tracking data were imported into R Studio (v3.6.1) and processed with the DLCAnalyzer package (Sturman et al., 2020). Furthermore, a previously trained supervised classifier (Sturman et al., 2020) was applied to quantify supported and unsupported rears in the OFT on a per-frame basis.

## Data availability

Sequencing data will be made available upon publication.

## Code availability

All code for analyzing sequencing experiments are available on https://github.com/ETHZ-INS/CRS_habituation.

## Author contribution

**Rebecca Waag:** conceived experiments, conducted experiments and molecular screens, data analysis, graphs & figures, interpreted results, wrote manuscript

**Lukas von Ziegler:** data analysis (transcriptomics)

**Emanuel Sonder:** data analysis (single-cell ATAC)

**Oliver Sturman:** help with data analysis (DLC behavior)

**Justine Leonardi:** practical help with experiments

**Selina Frei:** performed corticosterone ELISA

**Katharina Gapp:** supervision

**Pierre-Luc Germain:** data analysis (transcriptomics, ATAC, single-cell transcriptomics), conceived experiments, graphs & figures, interpreted results, provided funding, wrote manuscript

**Johannes Bohacek:** conceived experiments, interpreted results, provided resources and funding, wrote manuscript

All authors have read the manuscript.

## Acknowledgements

The lab of J.B. is supported by the ETH Zurich, ETH Project Grant ETH-20 19-1, SNSF Grant 310030_204372, the Basel Research Centre for Child Health (BRCCH), the Swiss 3R Competence Center, the Hochschulmedizin Zürich Flagship project STRESS, and ERA-NET NEURON - PROGRESS (SNSF: 31NE30_219119). E.S was supported by the ETH Project Grant ETH-25 20-2 to P.-L.G. We thank Julia Bode for maintaining the animal colony, Fabienne Rössler for help with data analysis, and Rosie Longster for experimental help. We thank the 3R Hub at the ETH Zurich for their support with behavior testing, and the staff of the EPIC for the excellent animal care and their service to our animal facility. Parts of figures were created using Biorender.com.

---

**Supplementary Figure 1:**
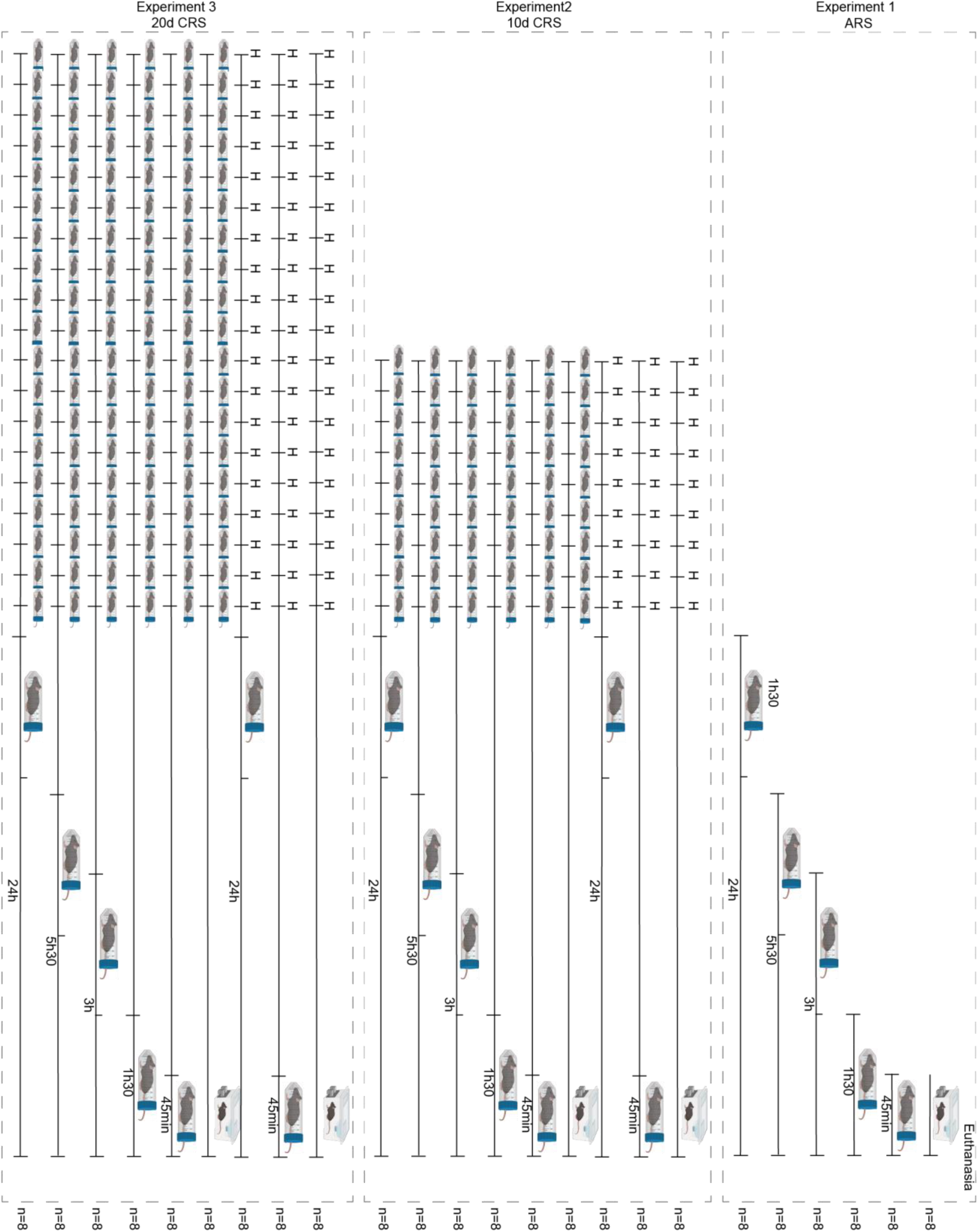
Experimental groups used for bulk transcriptional analysis. H = handling.

**Supplementary Figure 2:**
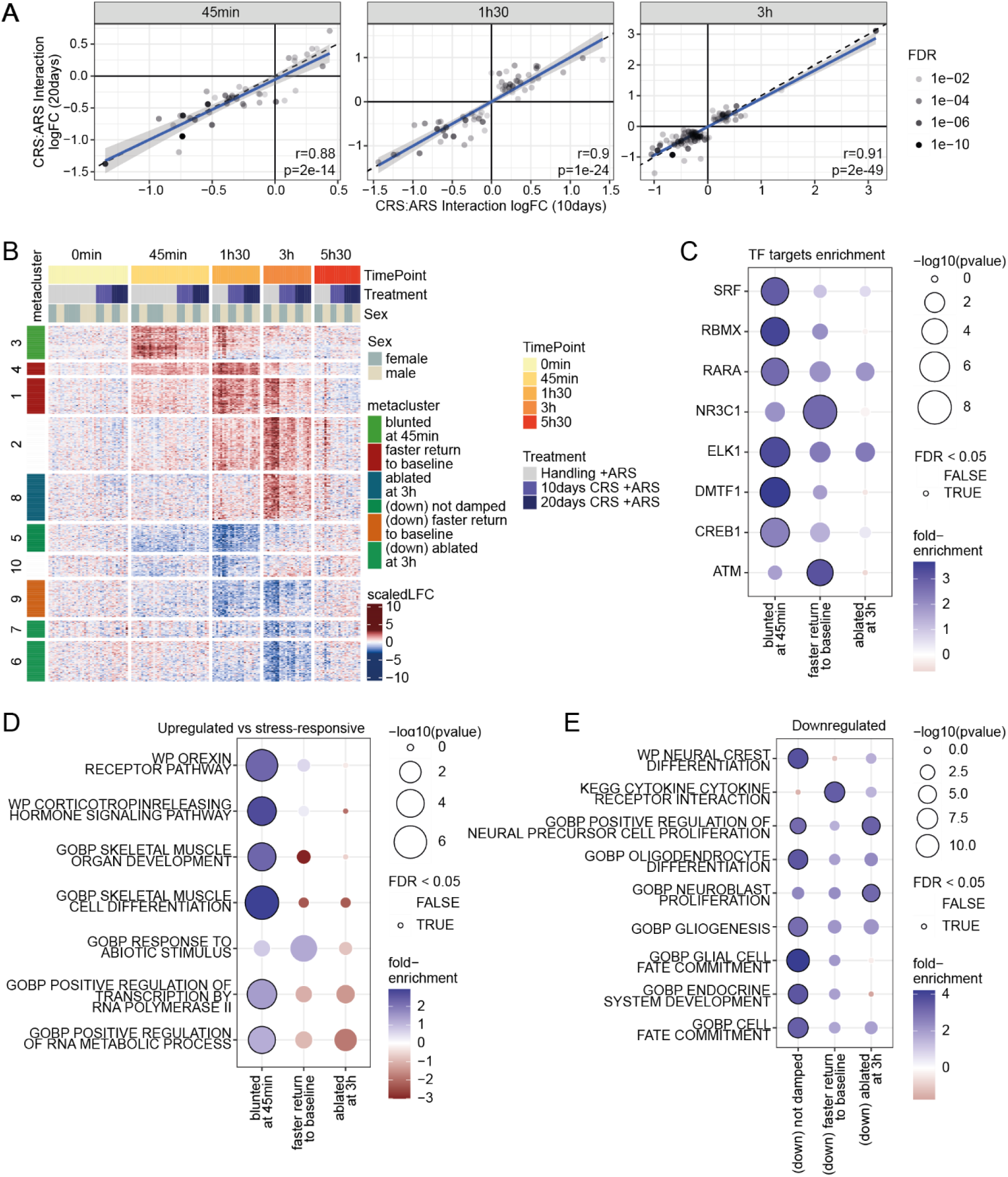
**A:** Comparison of the CRS:ARS interaction coefficients across the two treatment durations, for each ARS time point individually. **B:** Spectral clustering of the stress-responsive genes, highlighting the selected clusters. **C:** Transcription Factor target enrichment of the selected clusters. **D:** GO/Pathway enrichments of the clusters when compared to all stress-responsive genes. **E:** GO/Pathway enrichments of the downregulated clusters (compared to all tested genes).

**Supplementary Figure 3:**
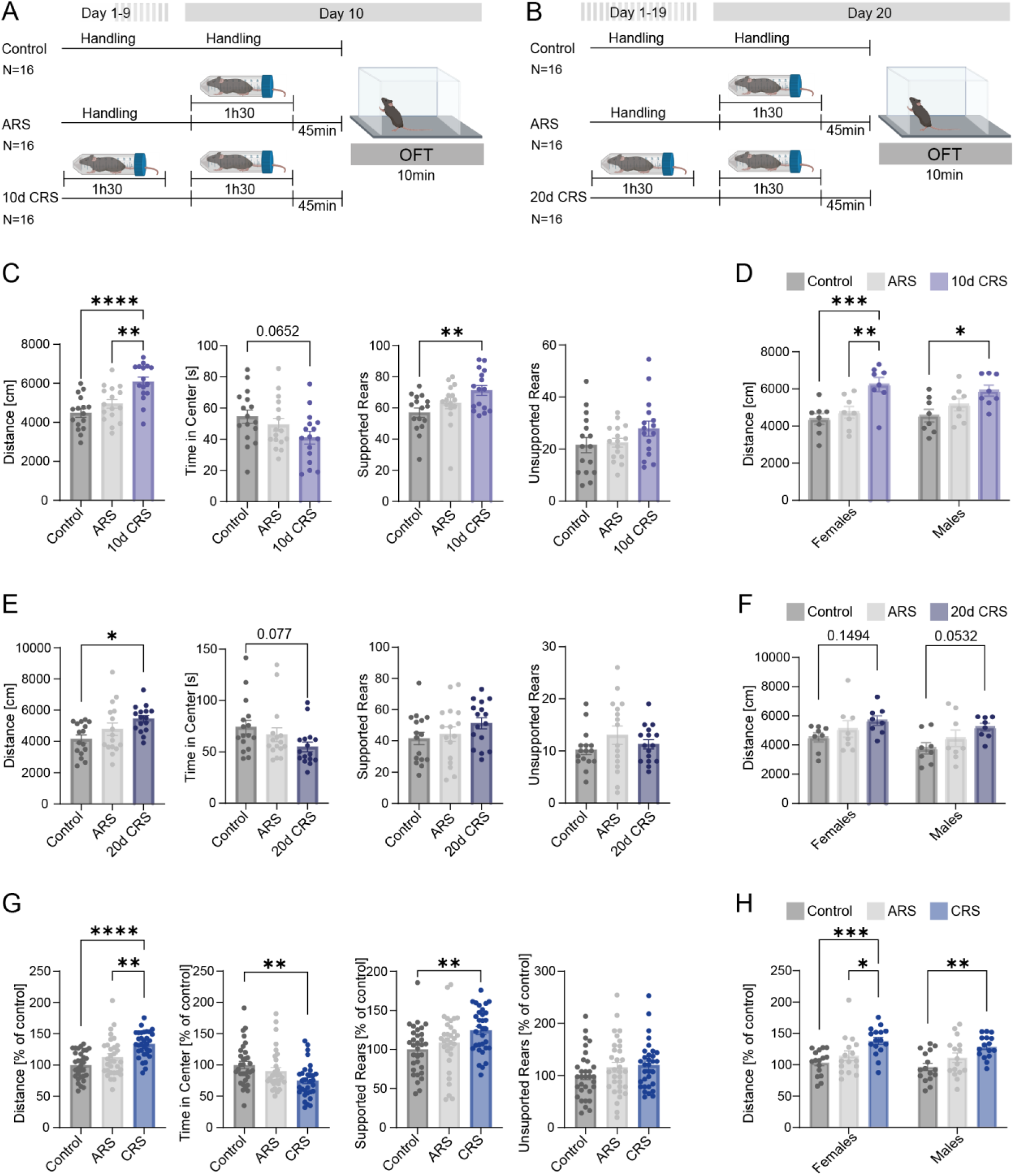
Effects of ARS and CRS on anxiety-like behavior in the open-field test (OFT) **A:** Experimental Design of 10d CRS experiment. **B**: Experimental Design of 20d CRS experiment. **C**: Behavioral phenotyping in the OFT after 10d CRS (1-way ANOVA, Tukey’s multiple comparison). **D**: Distance covered for female and male animals separately after 10d CRS (2-way ANOVA, Tukey’s multiple comparison). **E**: Behavioral phenotyping in the OFT after 20d CRS (1-way ANOVA, Tukey’s multiple comparison). **F**: Distance covered for female and male animals separately after 20d CRS (2-way ANOVA, Tukey’s multiple comparison). **G**: 10d and 20d CRS normalized to the respective control groups (1-way ANOVA, Tukey’s multiple comparison). **H**: Distance covered for female and male animals separately after 10d and 20d CRS (2-way ANOVA, Tukey’s multiple comparison). Data expressed as mean ± SEM. *adjusted p-value<0.05, **adjusted p-value<0.01, ***adjusted p-value<0.001, ****adjusted p-value<0.0001.

**Supplementary Figure 4:**
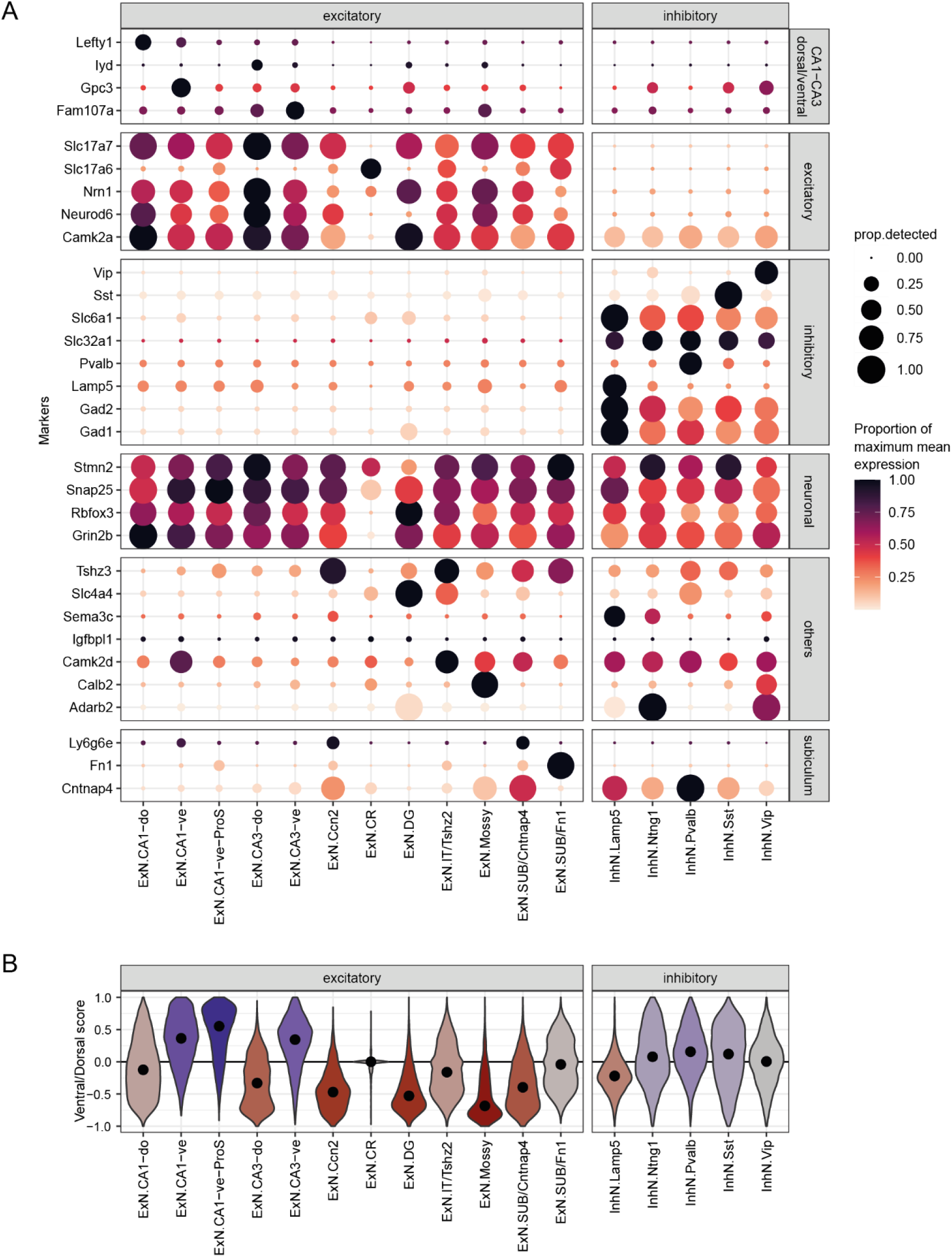
**A:** Expression of distinct sets of markers (bottom) across the clusters. **B:** Distributions of single-cell ventral/dorsal scores.

**Supplementary Figure 5:**
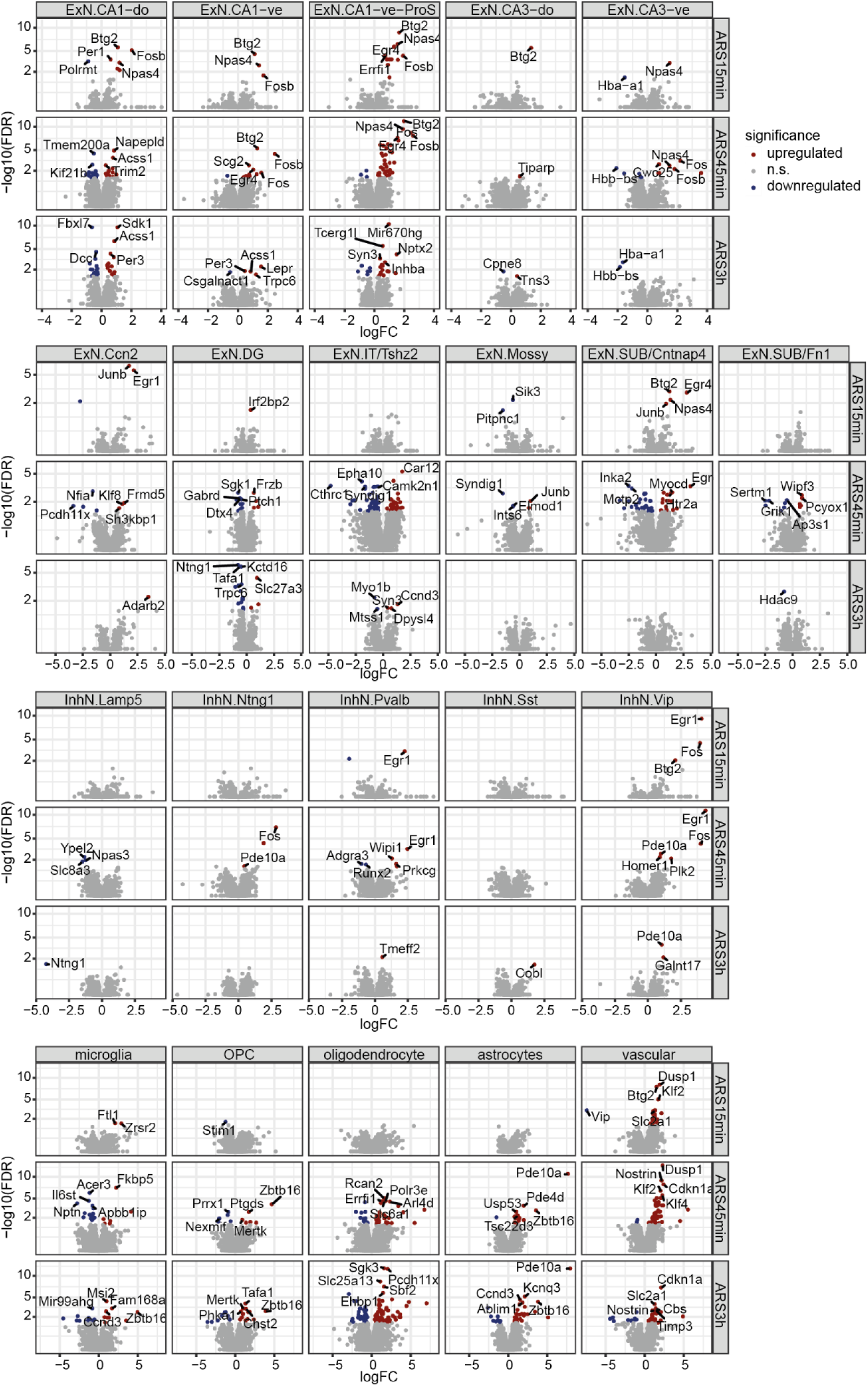
Volcano plot of the differential expression in each cell type at each time point upon ARS. Genes with FDR<0.05 are colored in red (increase) or blue (decrease), and the top significant genes in each facet are labeled.

**Supplementary Figure 6.**
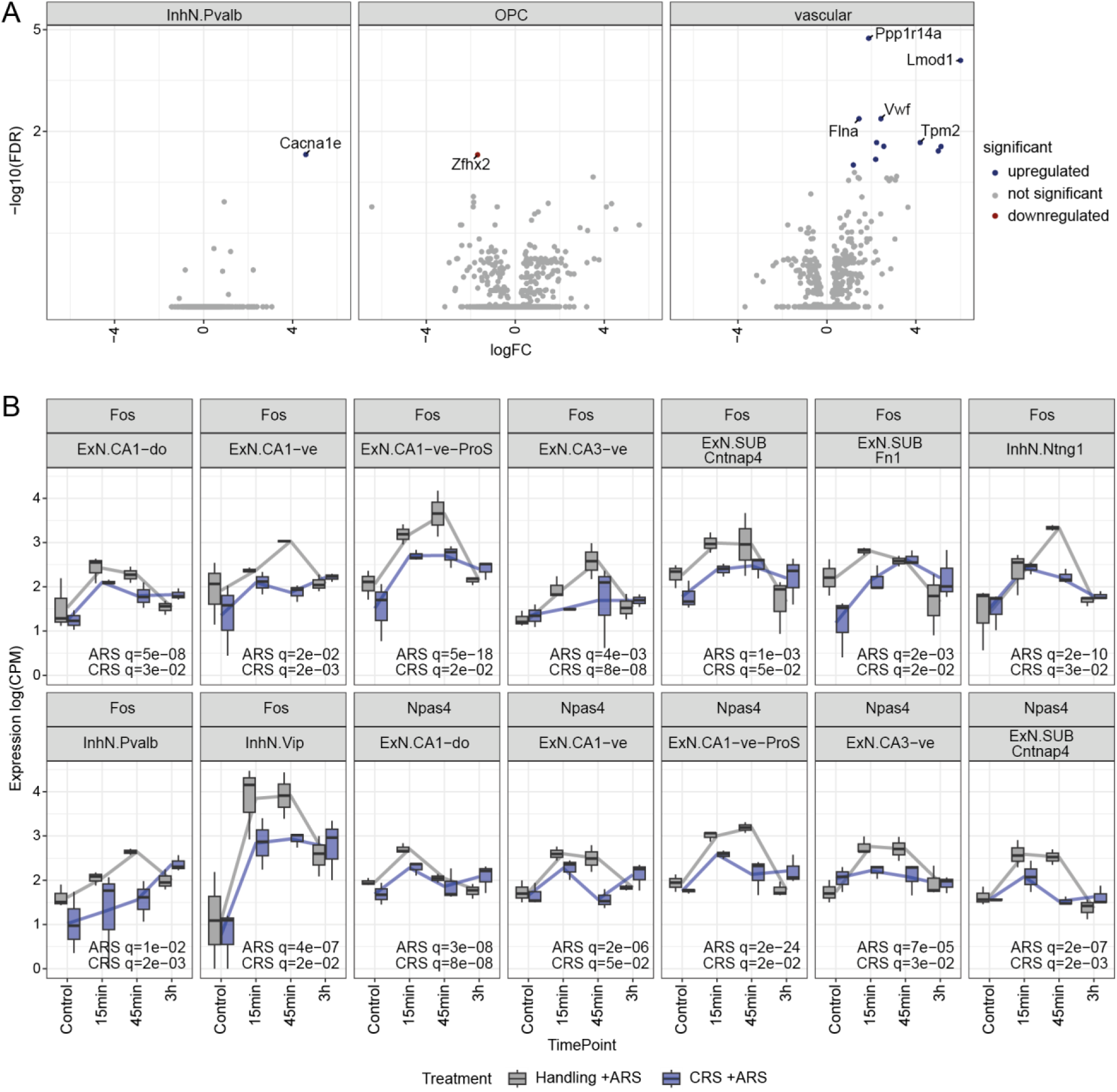
**A:** Volcano plots showing the significant changes in CRS at baseline. Genes with FDR<0.05 are colored in blue, and the top significant genes in each cell type are labeled. **B:** Expression of *Fos* and *Npas4* in the cell types in which they are significantly regulated by ARS. Reported significance values are from a likelihood ratio test, comparing the full model (ARS time points and CRS covariates, i.e. ~ARS*CRS) to a null model without the time points (ARS p-value) or without the CRS covariates (CRS p-value). Adjustment for multiple testing using Benjamini & Hochberg’s method. A significant CRS q-value means that CRS affects either the baseline gene expression or that the gene responds differently to ARS after CRS.

**Supplementary Figure 7:**
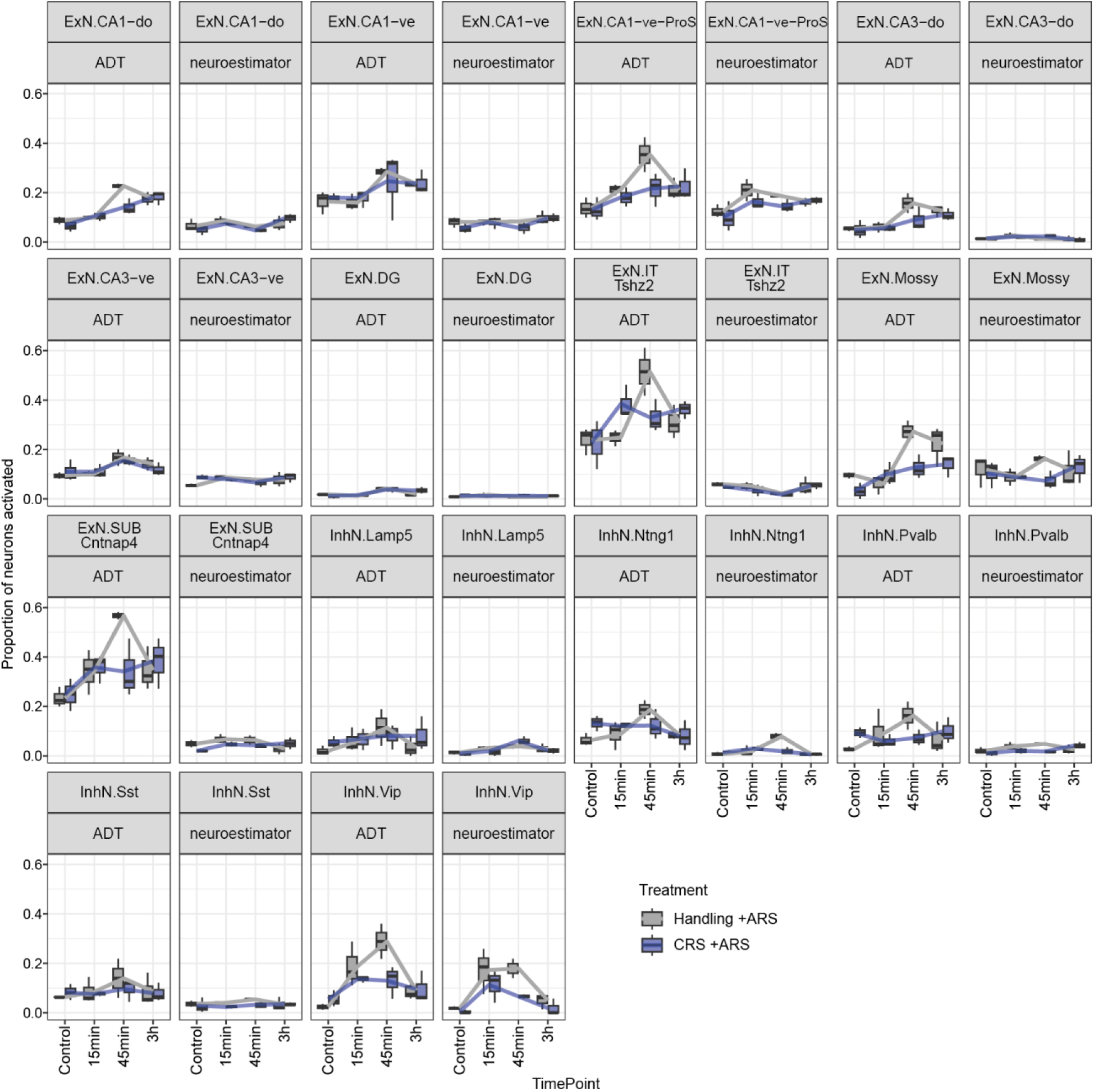
Comparison of the proportions of neurons activated based on the Activity-Dependent Transcription (ADT) score or on neuroestimator.

**Supplementary Figure 8:**
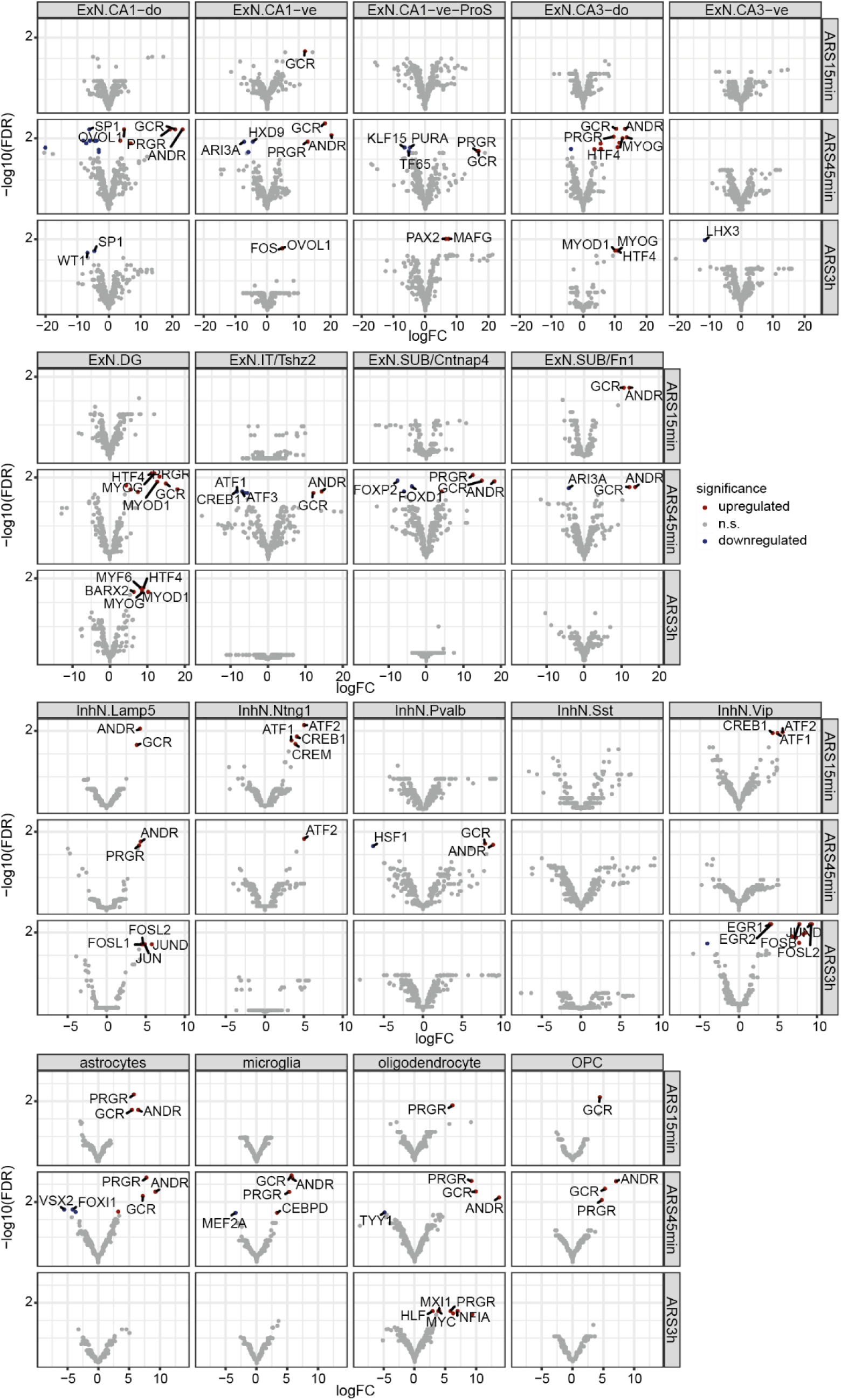
Volcano plots showing the significant changes in motif activity upon ARS. Motifs with FDR<0.05 are colored in blue, and the top significant ones in each facet are labeled.

**Supplementary Figure 9.**
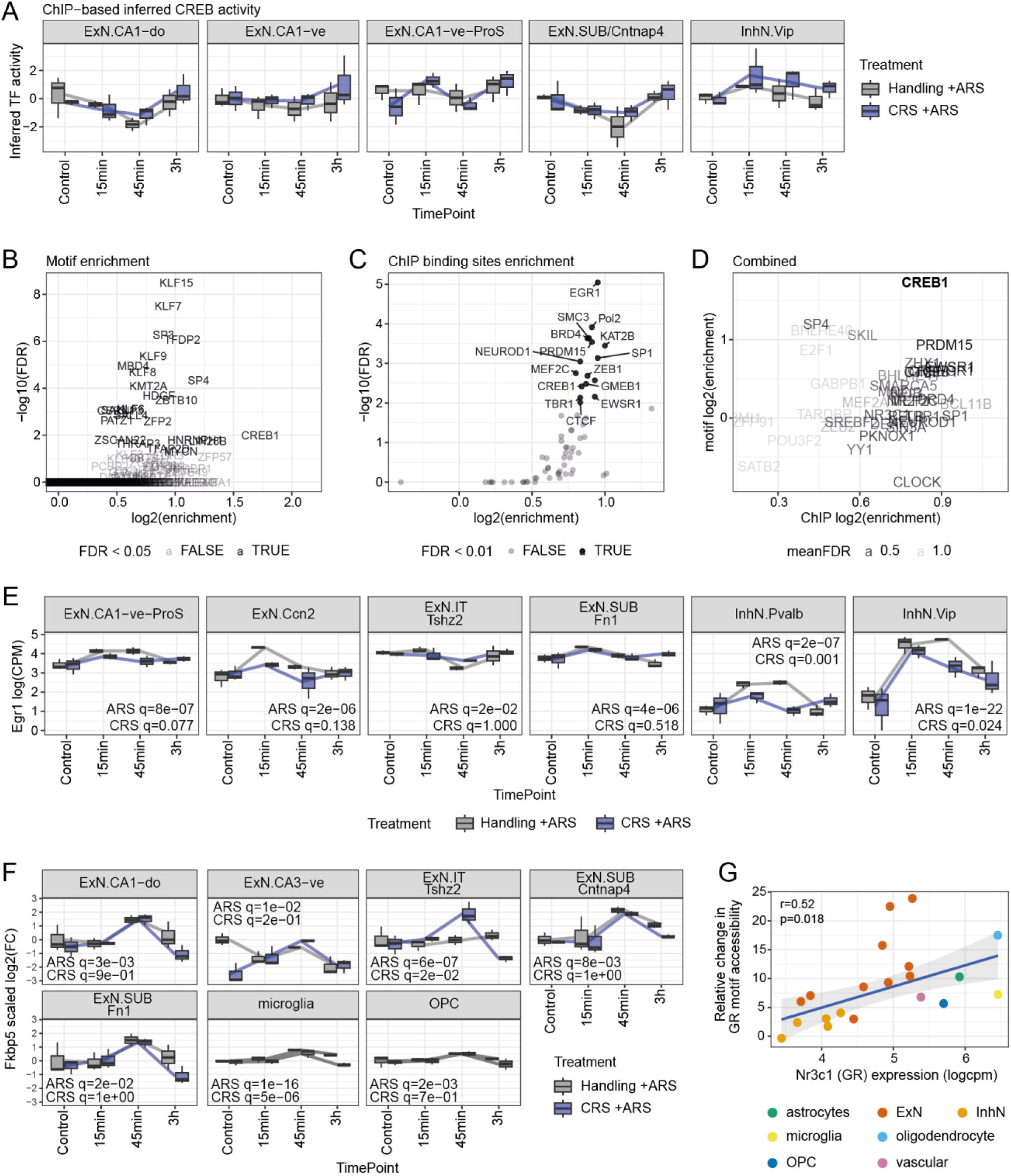
**A:** Inferred CREB1 activity based on the fastMLM method using the brainTF peaks. **B:** Motif enrichment at the neuronal peaks of genes responding early to ARS (i.e. significant at either 15min in the single-nucleus data or 45min in the spliced fraction of the bulk transcriptome). **C:** Enrichment in brainTF ChIP-seq peaks at the same peaks of ARS-responding genes. **D:** Combination of B and C, with CREB1 being the only TF with a mean FDR <0.05 in both motif- and ChIP-enrichment analyses. **E:** RNA expression of *Egr1* in the cell types in which it is statistically significant. **F:** Relative RNA levels of *Fkbp5* in the cell types in which it is statistically significant. **G:** The extent of increase in GR motif accessibility is correlated with the expression of the receptor.

